# Genomic analysis of *Ancylistes closterii*, an enigmatic alga parasitic fungus in the arthropod-associated Entomophthoromycotina

**DOI:** 10.1101/2023.12.22.573025

**Authors:** K Seto, TY James

**Affiliations:** Department of Ecology and Evolutionary Biology, University of Michigan, Ann Arbor, MI, 48109, USA; Faculty of Environment and Information Sciences, Yokohama National University, Yokohama, Kanagawa 240-8501, Japan

**Keywords:** comparative genomics, parasite, phylogenomics, zygomycete fungi

## Abstract

Recent advances in fungal genome sequencing have dramatically altered our understanding of the phylogeny and evolution of Fungi. However, there are still many poorly studied obligate parasitic or symbiotic fungi for which we lack any genomic information or knowledge of where they fit in the fungal phylogeny. *Ancylistes,* an endoparasite of desmid green algae, is such an understudied fungal genus. This genus has been taxonomically placed in the group of arthropod pathogens and saprobes, Entomophthoromycotina in Zoopagomycota. Understanding the phylogenetic position of *Ancylistes* provides insights into the nutritional evolution of Zoopagomycota, which is primarily composed of animal-associated fungi. In this study, we found and cultivated *Ancylistes closterii* with its host *Closterium* sp. and sequenced its genome to investigate its phylogenetic position and evolution. Phylogenetic analyses using rDNA and genome-scale datasets showed that *A. closterii* was sister to other Entomophthoromycotina fungi, confirming the taxonomic position of *Ancylistes*. Despite the ecological distinctiveness between *Ancylistes* and other Entomophthoromycotina fungi, our comparative genomic analyses revealed many shared traits of these fungi such as lineage-specific subtilases and hybrid histidine kinases. *Ancylistes* also possessed unique genes among Zoopagomycota fungi, such as plant cell wall degrading enzymes which could be important for infection of algae.

**Significance:** Improved taxon sampling is important for inferring a robust phylogeny of Fungi. However, there are still poorly studied obligate parasitic taxa whose DNA sequencing is challenging, especially in Zoopagomycota, one of the early diverging lineages of Fungi. This study focused on a long-neglected algal parasite, *Ancylistes closterii*, which belongs to the arthropod-associated group, Entomophthoromycotina. We rediscovered *A. closterii* and established a dual culture of fungus and its host alga, which enabled the first molecular analysis of this enigmatic parasite. Our results provide new insights into the nutritional evolution of primarily animal-associated Zoopagomycota.

## Introduction

Genomes provide the instructions organisms use for development and interactions with their environment. The recent revolution in sequencing capacity holds great promise for leveraging genomes to provide a deeper understanding of traits of many non-model species. These insights are particularly needed for microbial taxa, such as prokaryotes and fungi, where many of the lineages have not been observed or cultured. Unculturable taxa pose problems both for sequencing and for morphological analyses, however, for parasitic taxa, some can be maintained on their host, and the use of dual host-parasite cultures has been pivotal for genomic analysis of these parasites (Galindo et al. 2021; James et al. 2013; Mikhailov et al. 2022). Many of the early diverging lineages of fungi, such as Chytridiomycota and Microsporidia, are considered obligate parasites, and the need for both single-cell genomics and dual host-parasite cultures is pronounced. Investigation of the genomes of these parasites is needed to better understand their metabolism and symbioses with their hosts and the evolution of these characteristics throughout all fungi (Rahimlou et al. in press).

Zygomycete fungi, which were formerly included in a single phylum Zygomycota, are important lineages to investigate the early evolution of terrestrial fungi because they represent intermediate branches between zoosporic lineages and the highly successful and species-rich lineage, Dikarya (Ascomycota and Basidiomycota) (Spatafora et al. 2017). Reconstruction of a robust phylogeny is essential for evolutionary biology, however, phylogenetic analysis of zygomycete fungi has been challenging due to either poor lineage sampling or low number of marker genes employed. In many phylogenetic analyses of rDNA genes and/or a few protein-coding genes, Zygomycota was not recovered as a monophyletic group and separated into several clades (James et al. 2006; White et al. 2006), and, moreover, the relationships between clades were poorly resolved. During the last decade, phylogenomic approaches, which utilize hundreds to thousands of genes retrieved from genomic or transcriptomic data, were applied to zygomycete fungi and improved phylogenetic resolution (Spatafora et al. 2016; Chang et al. 2019; Davis et al. 2019; Wang et al. 2020; Chang et al. 2022; Reynolds et al. 2023). The results supported a classification that divided zygomycete fungi into two phyla both of which include three subphyla: Mucoromycota (Glomeromycotina, Mortierellomycotina, Mucoromycotina) sister to Dikarya, and Zoopagomycota (Entomophthoromycotina, Kickxellomycotina, Zoopagomycotina) sister to Dikarya + Mucoromycota (Spatafora et al. 2016). Although the availability of genome data of zygomycete fungi is increasing, it tends to be biased toward Mucoromycota taxa, in which Mucorales (Mucoromycotina) taxa are especially rich in their genomic data (Gryganskyi et al. 2023; Wang et al. 2023). This taxonomic bias could be because they are generally saprotrophic and thus easy to be cultured, and some of them are causal agents of mucormycosis (Soare et al. 2020). In contrast, many lineages of Zoopagomycota are represented by obligate parasitic taxa (Benny et al. 2014) whose genomic studies are often challenging. The difficulties of genome sequencing of parasitic taxa were partially overcome by new techniques such as single-cell genomics (Ahrendt et al. 2018; Davis et al. 2019). However, in Zoopagomycota, there are still poorly studied parasitic taxa that entirely lack DNA sequence data (Wijayawardene et al. 2018). One example is the neglected genus *Ancylistes*, which was examined in this study.

*Ancylistes* is an endoparasite of desmid green algae, *Closterium* and *Netrium* (Pfitzer 1872; Couch 1949). It develops in the host cell as hyphae which are segmented by septation in maturity, and each segment produces external hyphae extending outside of the host cell. A new infection occurs when external hyphae touch a new host cell. Although this fungus produces resting spores sexually, no asexual spores were initially recorded (Pfitzer 1872). This enigmatic fungus was placed in the order Ancylistales along with some endoparasitic oomycetes such as *Lagenidium* and *Myzocytium* (Schröter 1897). However, Berdan (1938) revealed that *Ancylistes* produces conidia forcibly discharged from conidiophores at the tip of external hyphae, which is strikingly similar to Entomophthorales. Based on this observation, *Ancylistes* was removed from Ancylistales and transferred to Entomophthorales. Entomophthorales (currently Entomophthoromycotina in Zoopagomycota) is well known as a group of arthropod pathogens but also includes some saprotrophs, and is characterized by coenocytic vegetative cells, production of forcibly discharged primary and secondary conidia, and zygospore formation with some exceptions (Gryganskyi et al. 2013). *Ancylistes* has been taxonomically placed in the family Ancylistaceae (Entomophthorales, Entomophthoromycetes) along with the genus *Conidiobolus* which includes primarily saprotrophic taxa and also some opportunistic human pathogens (Humber 1989, 2012). However, molecular phylogenetic analyses have shown that *Conidiobolus* was separated into several paraphyletic lineages sister to the arthropod pathogenic lineage, Entomophthoraceae (Gryganskyi et al. 2012, 2013), indicating the necessity of revision of Ancylistaceae. Recently, *Conidiobolus* sensu lato (s.l.) was separated into five genera, *Azygosporus*, *Capillidium*, *Neoconidiobolus*, *Microconidiobolus*, and *Conidiobolus* sensu stricto (s.s.) (Nie et al. 2020; Cai et al. 2021). These genera were removed from Ancylistaceae and accommodated in newly established families, Capillidiaceae (*Capillidium*), Conidiobolaceae (*Azygosporus*, *Conidiobolus,* and *Microconidiobolus*), Neoconidiobolaceae (*Neoconidiobolus*) although the phylogeny of *Ancylistes* has never been examined (Gryganskyi et al. 2022). DNA sequencing of *Ancylistes* is pivotal to justify the recent taxonomic revisions of Ancylistaceae. Furthermore, the phylogenetic placement of *Ancylistes* is also important to examine nutritional evolution in the group because members of Entomophthoromycotina as well as entirely Zoopagomycota are primarily associated with animals (Spatafora et al. 2016).

In this study, we rediscovered *Ancylistes closterii* from a pond in Michigan, USA, and cultivated it as a dual culture (strain KS117) with its host, *Closterium* sp. By utilizing this culture, we observed morphology in detail, sequenced its genome, and revealed the phylogenetic position of the genus for the first time. We then used comparative genomics to define the shared and unique traits of *Ancylistes* among Zoopagomycota lineages.

## Results

### Morphology of *A. closterii*

Based on morphological characters (Figs. 1, 2), strain KS117 was identified as *A. closterii*, the type species of the genus (Pfitzer 1872; Berdan 1938). The fungus developed as hyphae in the host (*Closterium* sp.) cell and consumed the host cytoplasm (Fig. 1A). The hyphae are divided into several segments by septation (Fig. 1B), and one hypha emerged from each segment and extended to the outside of the host cell (Fig. 1C–E). The external hyphae extended by pushing their cytoplasm towards the tip and delimiting by septation (Fig. 1F–I). The external hyphae reached a new host cell and invaded via an appressorium (Fig. 1J). When the fungus was on the surface of water, external hyphae formed conidiophores to produce forcibly discharged conidia (Fig. 2A–C). Secondary conidia formation and germination of conidia were observed (Fig. 2D–G). Under dark conditions, the fungus produced resting spores (Fig. 2H, I) by sexual reproduction in which fusion of two hyphae occurred (Fig. 2J–L). Observations on stained nuclei showed that conidia, external hyphae, and endobiotic thalli were multinucleated (Fig. S1).

**Fig. 1.**
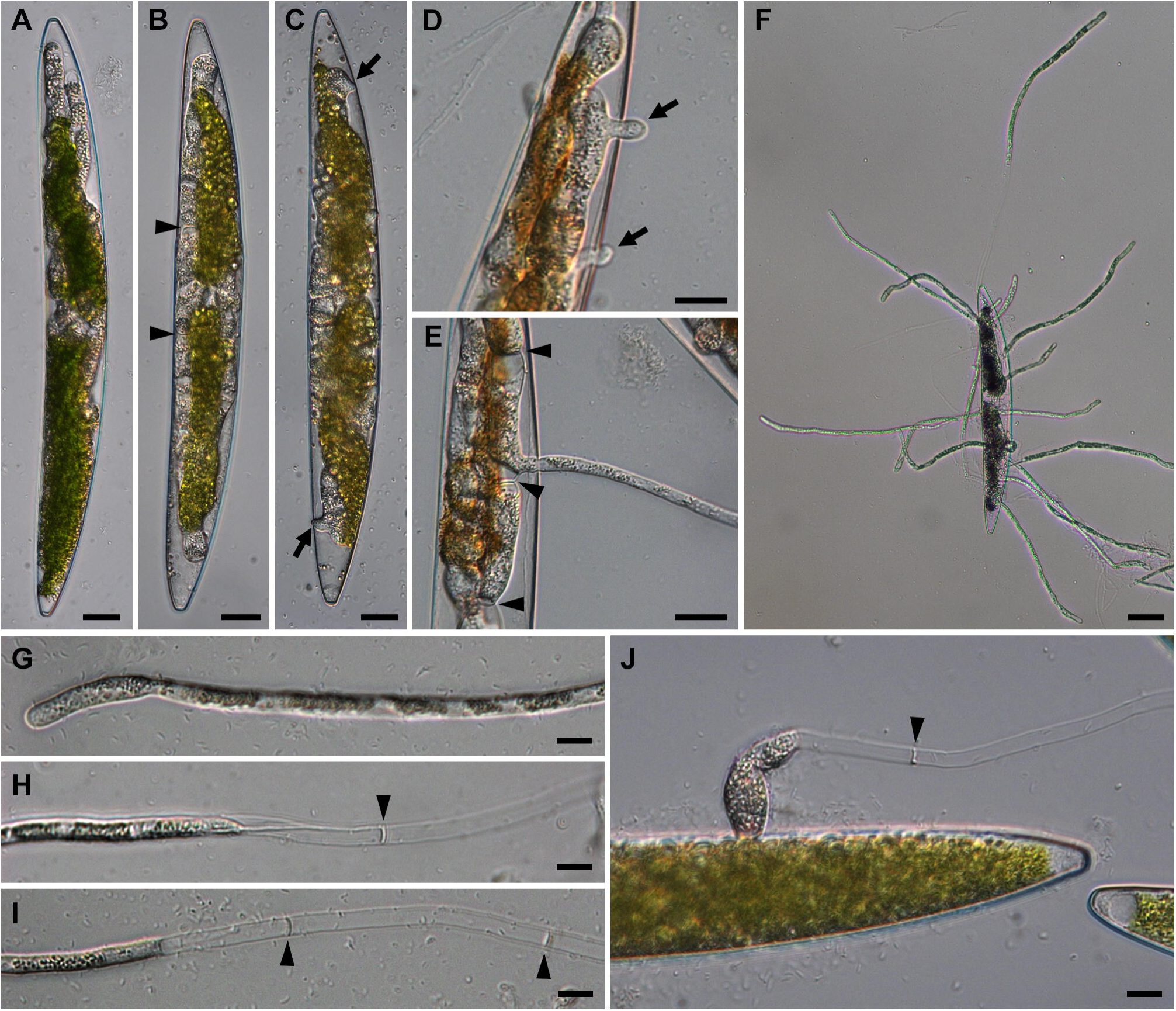
Development of *Ancylistes closterii* parasitic on *Closterium* sp. (*A*) Young parasite hyphae developing in a host cell. (*B*) Segmentation of parasite hyphae by septation (arrowheads). (*C*–*E*) Development of external hyphae (arrows). Panel *E* shows that one segment delimited by septations (arrowheads) produces one external hyphae. (*F*) Multiple external hyphae extending from a dead host cell. (*G*–*I*) Development of external hyphae. Panels *H* and *I* show that cytoplasm moves forward and that hyphae are delimited by regular septation (arrowheads). (*J*) New infection via appressorium at the hyphal tip delimited by septation (arrowhead). Bars = 20 µm (*A*–*E*), 50 µm (*F*), 10 µm (*G*–*J*).

**Fig. 2.**
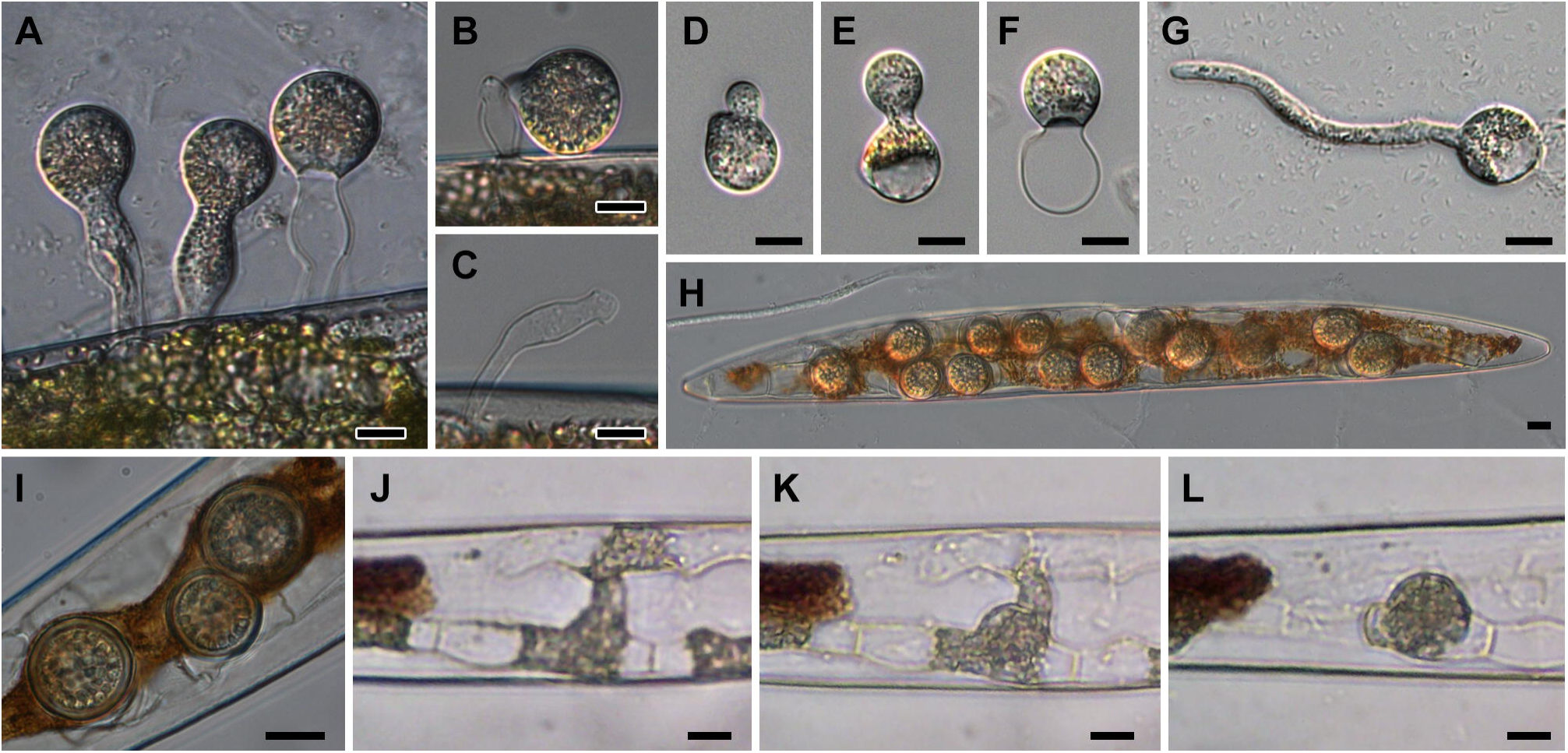
Formation of conidia and resting spores of *Ancylistes closterii*. (*A*) Conidia formed on conidiophores produced at the tips of external hyphae. (*B*) Detached conidium. (*C*) Conidiophore after conidium discharge. (*D*–*F*) Formation of secondary conidia. (*G*) Germinated conidium. (*H*, *I*) Resting spores in a host cell. (*J*–*K*) Time-lapse images of resting spore formation. Panels *I* and *K* are 1.5 and 3 hours after *J*, respectively. Bars = 10 µm.

### Genome assembly and phylogeny of *A. closterii*

The genome assembly of *A. closterii* comprised 2,121 scaffolds of total length 111.95 Mb (N50 = 125,571) with a 22% GC content. A BUSCO v4 (Manni et al. 2021) analysis using the fungi_odb10 database showed 80.5% (604 out of 758 genes) completeness of the genome. In total, 11,779 protein-coding genes were predicted. The *k*-mer and allele frequency analyses (Amses et al. 2022) indicated that the genome of *A. closterii* was haploid or homozygous (Fig. S2).

To investigate the phylogenetic position of *A. closterii*, we performed phylogenetic analyses using the 18S-28S rDNA and genome-scale datasets. Since rDNA contigs were fragmented in the aforementioned genome assembly, a targeted assembling of the rDNA operon was conducted using GetOrganelle (Jin et al. 2020). A maximum likelihood (ML) analysis on the concatenated dataset of 18S and 28S rDNA (Table S1) showed that *A. closterii* was placed in Entomophthoromycotina along with Entomophthoraceae and *Conidiobolus* s.l. (Fig. S3). The four major clades of *Conidiobolus* s.l. were recovered, which correspond to the families Batkoaceae, Capillidiaceae, Conidiobolaceae, and Neoconidiobolaceae (Gryganskyi et al. 2022). The genus *Microconidiobolus* was not monophyletic in our analysis. Phylogenetic relationships between lineages in Entomophthoromycotina were poorly supported, probably due to highly divergent sequences of rDNA regions in these fungi.

To infer a robust phylogeny, phylogenomic analyses using 49 fungal genomes (Table S2) were performed. A supermatrix comprising 185,578 amino acids (aa) of 452 genes in the fungi_odb10 database was prepared for these analyses. Preliminary ML analyses with site-homogeneous (LG+F+R10) and site-heterogeneous (LG+C20+F+G with PMSF method) (Wang et al. 2018) models showed incongruence of tree topologies (Figs. S4, S5). In the tree of the site-homogeneous model, all zygomycete fungi (Mucoromycota + Zoopagomycota) were monophyletic and *Basidiobolus* was sister to Mucoromycota (Fig. S4). In contrast, in the site-heterogeneous model tree (Fig. S5), zygomycete fungi were paraphyletic and *Basidiobolus* was sister to other Zoopagomycota. We hypothesized that fast-evolving sites caused incorrect phylogenetic signals in the site-homogeneous model analysis as observed in the previous analyses (Brown et al. 2013; Kang et al. 2017). The effect of fast-evolving sites was assessed by removing the fastest-evolving sites in a stepwise fashion (10,000 aa per step) and recording ultrafast bootstrap (UFBoot) values of the interested nodes. The result showed that the placement of *Basidiobolus* was affected by fast-evolving sites (Fig. S6). The UFBoot value of “*Basidiobolus* + Mucoromycota” and the one for monophyly of all zygomycete fungi (Mucoromycota + Zoopagomycota + *Basidiobolus*) were identical, and they decreased in tandem by removing fast-evolving sites. In its place, the support of “*Basidiobolus* + other Zoopagomycota” increased. When the 40,000 fastest-evolving sites were removed, the same tree topology as the site-heterogeneous model tree was obtained. To minimize phylogenetic errors, the supermatrix of 135,578 aa formed after removing the 50,000 fastest-evolving sites was used for the following analyses. Two site-heterogeneous model analyses, ML analysis with LG+C60+F+G+PMSF model (Fig. 3) and Bayesian analysis with CAT-Poisson+G4 model (Fig. S7), recovered the same topology except for the monophyly of Mucoromycota, in the latter of which it was paraphyletic. *Ancylistes closterii* was sister to all other Entomophthoromycotina taxa, which was fully supported by both analyses. In Entomophthoromycotina, *Conidiobolus* s.l. was paraphyletic, the clade of “*Capillidium heterosporum* + *Neoconidiobolus thromboides*” was sister to Entomophthoraceae (*Entomophthora muscae* + *Zoophthora radicans*) and *Conidiobolus coronatus* was placed outside of these taxa. In Zoopagomycota, Zoopagomycotina was sister to “Entomophthoromycotina + Kickxellomycotina”, and two *Basidiobolus* species were sister to all other Zoopagomycota lineages. Although *Basidiobolus* is morphologically classified as Entomophthoromycotina (Humber 2012), its phylogenetic position is controversial (Li et al. 2021). In this study, we place *Basidiobolus* at an independent position in Zoopagomycota without giving a higher taxonomic rank. The approximately unbiased (AU) test (Shimodaira 2002) showed that alternative tree topologies such as placements of *Basidiobolus*, branching orders of *Conidiobolus* s.l., and monophyly of *Conidiobolus* s.l. and Ancylistaceae s.l. were significantly rejected (Table S3).

**Fig. 3.**
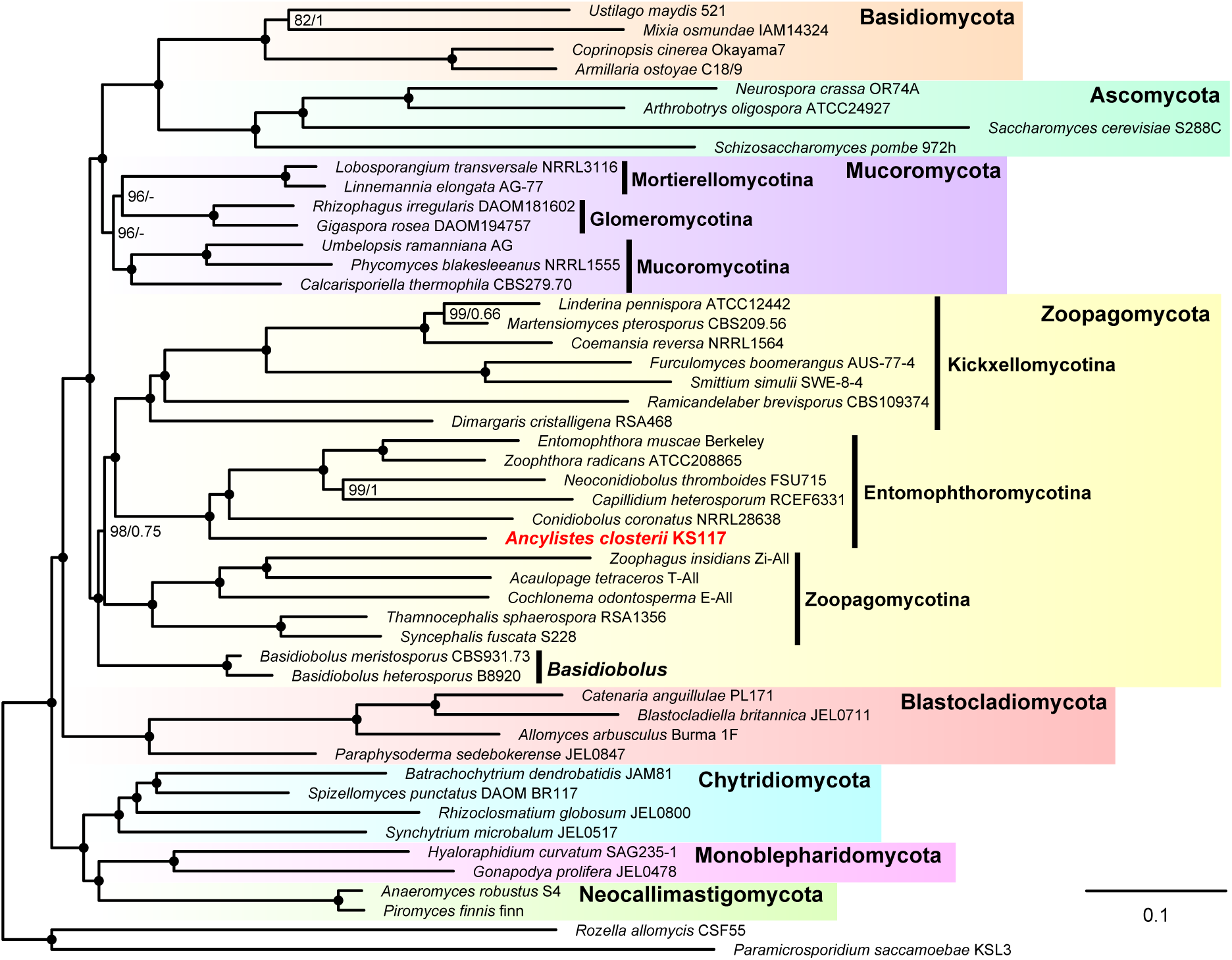
Maximum likelihood (ML) tree reconstructed with a concatenated dataset after removal of the 50,000 fastest evolving sites (135,578 amino acids of 452 genes). The tree was inferred with the LG+C60+F+G+PMSF model using IQ-TREE2, and support values of nodes were calculated with the standard bootstrapping analysis (100 replicates). Values next to nodes indicate ML bootstrap values (left) and Bayesian posterior probabilities (right) inferred with PhyloBayes MPI v1.8 (CAT-Poisson+G4 model). Nodes fully supported by both bootstrap value (=100) and posterior probability (= 1.00) were indicated by black circles.

### Genome comparison between *A. closterii* and other zygomycete fungi

To investigate the uniqueness of *A. closterii* in genomic aspects, we compared *A. closterii* with other zygomycete fungi. First, we investigated general differences in genomes between *A. closterii* and other Entomophthoromycotina fungi by the gene ontology (GO) enrichment analyses. We also focused on specific genes: plant cell wall degrading enzymes (PCWDEs), subtilases, hybrid histidine kinases (HHKs), and elongation factor 1α (EF1α)/elongation factor-like (EFL) genes. For these analyses, 27 zygomycete proteomes used for phylogenomic analyses (Mucoromycota and Zoopagomycota in Table S2) were examined.

The enrichment analyses showed that 70 and 26 GO terms were significantly (p-value adjusted by the Benjamini & Hochberg False Discovery Rate correction < 0.01) over- and underrepresented, respectively, in *A. closterii* compared to other Entomophthoromycotina taxa (Table S4). The hierarchical organizations of the over- and underrepresented GO terms in the “Molecular Function” category are shown in Figure S8. In these, two overrepresented terms (GO:0008810 [cellulase activity] and GO:0030570 [pectate lyase activity]) may reflect the algal-associated nature of *A. closterii* in contrast to insect pathogenic or saprotrophic lifestyles of other Entomophthoromycotina taxa (Fig. 4A). The GO terms related to peptidase activities, especially the ones of serine-type (e.g., GO:0004252 [serine-type endopeptidase activity]) were underrepresented in *A. closterii* (Fig. 4B). This reflects a lower number of serine proteases in *A. closterii*, and a much higher number in two insect pathogenic species, *E. muscae* and *Z. radicans* (Fig. 4C, total gene counts are based on the GO:0004252). Three GO terms, GO:0020037 (heme binding), GO:0016705 (oxidoreductase activity, acting on paired donors, with incorporation or reduction of molecular oxygen), and GO:0004497 (monooxygenase activity) were also underrepresented in *A. closterii*. In the taxa of *Conidiobolus* s.l. and Entomophthoraceae, genes with these GO term annotations were mostly represented by cytochrome P450 monooxygenases (Fig. 4D, Pfam annotation: PF00067). Cytochrome P450 monooxygenases are known to be highly expanded in *Conidiobolus* s.l. lineages (Ngwenya et al. 2018; Wang et al. 2020), and our results indicate that these expansions have not occurred in *A. closterii*.

**Fig. 4.**
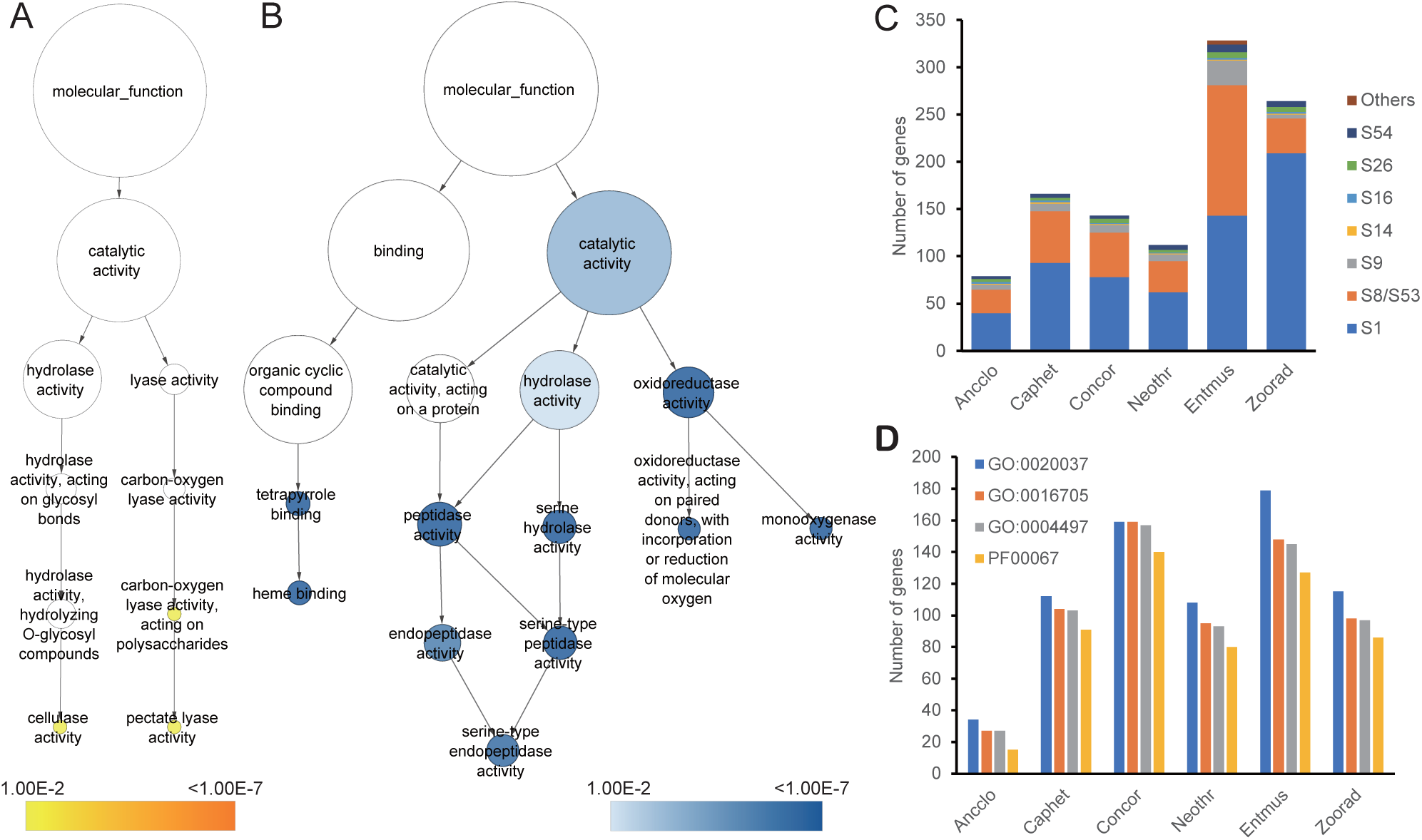
Genomic comparison between *Ancylistes closterii* and other Entomophthoromycotina taxa. (*A*, *B*) Hierarchical organizations of selected overrepresented (*A*) and underrepresented (*B*) gene ontology (GO) terms in *A. closterii* compared to other Entomophthoromycotina taxa. Colored circles indicate significantly overrepresented (yellow to orange) and underrepresented (light blue to dark blue) GO terms. (*C*) Number of genes annotated with the term GO:0004252 (serine-type endopeptidase activity) in each Entomophthoromycotina taxa. Subfamily compositions of serine-type peptidases are indicated by colors. (*D*) Number of genes annotated with the terms GO:0020037 (heme binding), GO:0016705 (oxidoreductase activity, acting on paired donors, with incorporation or reduction of molecular oxygen), and GO:0004497 (monooxygenase activity), and with the Pfam PF00067 (cytochrome P450 monooxygenases) in each Entomophthoromycotina taxa.

PCWDEs are a subset of carbohydrate-active enzymes (CAZymes) involved in the degradation of polysaccharides included in plant cell walls. In the zygomycete genomes, 23 PCWDEs degrading cellulose (n=7), hemicellulose (n=8), and pectin (n=8) were found (Fig. 5). *Ancylistes closterii* possessed six enzyme families degrading cellulose (GH3 and GH45), hemicellulose (CE1 and GH44), and pectin (GH35 and PL3_2). While the CE1, GH3, and GH35 genes were also found in many other zygomycete fungi, the GH44, GH45, and PL3_2 were uniquely expanded in *A. closterii* (Fig. 5). In the examined zygomycete fungi, only *A. closterii* possessed one GH44 gene, which is known as xyloglucanase and found in Basidiomycota fungi and bacteria (Sun et al. 2022). The gene tree showed that the GH44 gene of *A. closterii* was related to genes from Basidiomycota along with some genes of Chytridiomycota and Monoblepharidomycota (Fig. S9). Four copies of the GH45 genes, which are known as endoglucanase hydrolyzing β-1,4 glucans (Payne et al. 2015), were found in *A. closterii* but also in *Basidiobolus heterosporus* and two Mucoromycotina taxa (Fig. 5). In the gene tree (Fig. S10), the genes from *A. closterii* and the ones from *B. heterosporus* + Mucoromycotina taxa were separated, indicating their independent origin. Similarly, the PL3_2, which functions as pectate lyase (Garron & Cygler 2010), were found in *A. closterii* and *Linnemannia elongata* (Mortierellomycotina), but they were phylogenetically distinct in the gene tree (Fig. S11).

**Fig. 5.**
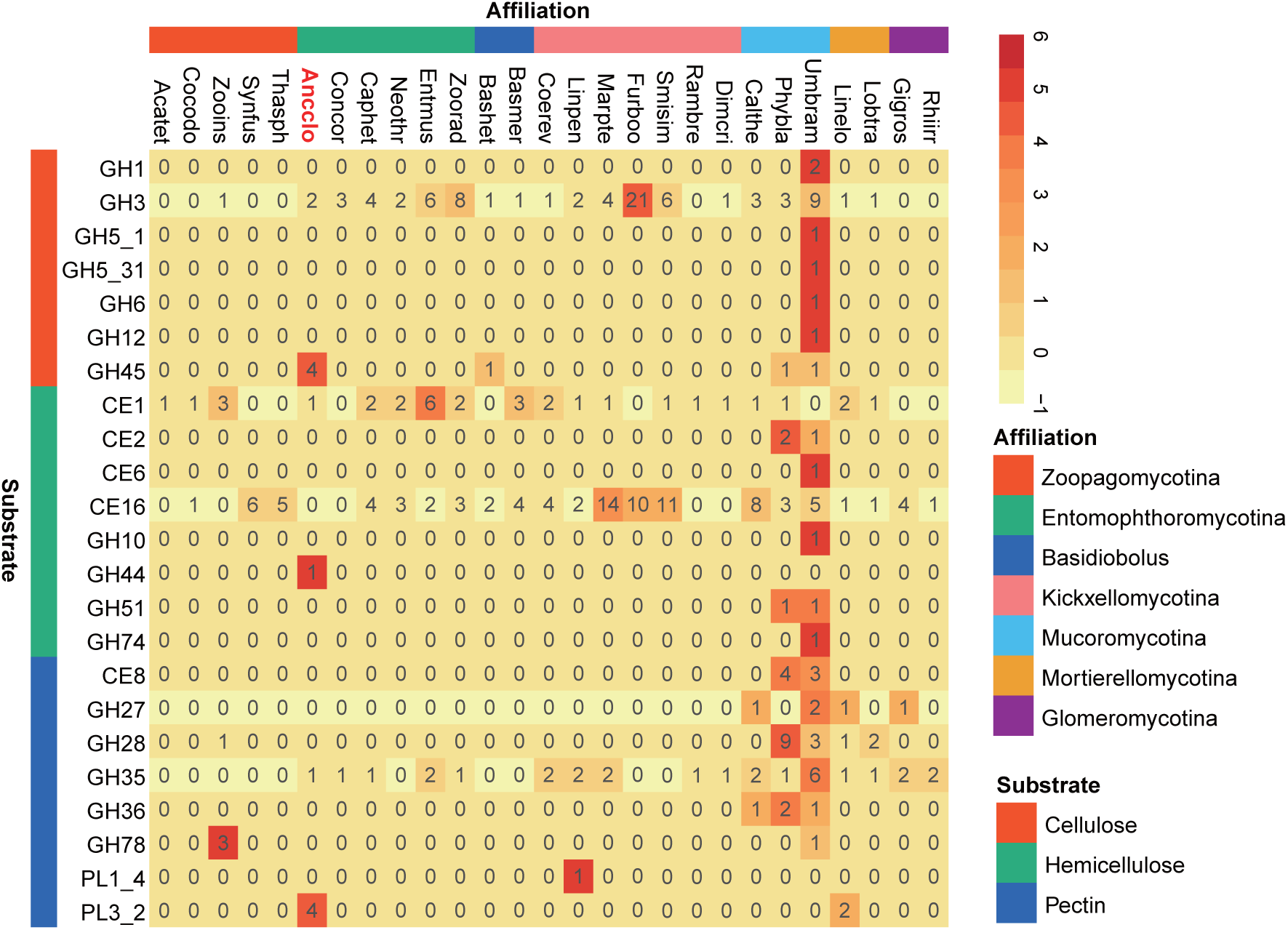
Heatmap showing plant cell wall degrading enzymes repertoire of zygomycete fungi. Number in each cell are raw count of each enzyme in each genome. Color in each cell indicates a z-score showing the relative abundance of each enzyme among the compared taxa. Species: Acatet, *Acaulopage tetraceros*; Ancclo, *Ancylistes closterii*; Bashet, *Basidiobolus heterosporus*; Basmer, *Basidiobolus meristosporus*; Calthe, *Calcarisporiella thermophila*; Caphet, *Capillidium heterosporum*; Cocodo, *Cochlonema odontosperma*; Coerev, *Coemansia reversa*; Concor, *Conidiobolus coronatus*; Dimcri, *Dimargaris cristalligena*; Entmus, *Entomophthora muscae*; Furboo, *Furculomyces boomerangus*; Gigros, *Gigaspora rosea*; Linelo, *Linnemannia elongata*; Linpen, *Linderina pennispora*; Lobtra, *Lobosporangium transversale*; Marpte, *Martensiomyces pterosporus*; Neothr, *Neoconidiobolus thromboides*; Phybla, *Phycomyces blakesleeanus*; Rambre, *Ramicandelaber brevisporus*; Rhiirr, *Rhizophagus irregularis*; Smisim, *Smittium simulii*; Synfus, *Syncephalis fuscata*; Thasph, *Thamnocephalis sphaerospora*; Umbram, *Umbelopsis ramanniana*; Zooins, *Zoophagus insidians*; Zoorad, *Zoophthora radicans*. Enzymes: GH, glycosyl hydrolase; CE, carbohydrate esterase; PL, polysaccharide lyase.

Subtilases (subtilisin-like serine proteases) are one of the superfamilies of serine peptidases (Siezen et al. 2007). Some subtilases are known to function as virulence factors in insect pathogenic fungi (Donatti et al. 2008; Fang et al. 2009). Recent studies showed two new subgroups of subtilases specific to Zoopagomycota lineages (Ahrendt et al. 2018; Arnesen et al. 2018). Investigation of subtilases based on the annotation of Pfam domain PF00082 (Subtilase family) showed 25 subtilases in *A. closterii*. A clustering analysis based on the sequence similarities of proteins showed that subtilases in *A. closterii* were categorized into five clusters (Fig. 6): 16 in proteinase K-like subtilases, 2 in kexins, 1 in “New 4 (Li et al. 2017)”, 1 in “Zoopagomycota cluster (Ahrendt et al. 2018)”. The other 5 genes were placed in a cluster exclusively composed of Entomophthoromycotina genes (except for one gene from Zoopagomycotina), which corresponds to the formerly reported Entomophthoromycotina-specific subtilases (Arnesen et al. 2018).

**Fig. 6.**
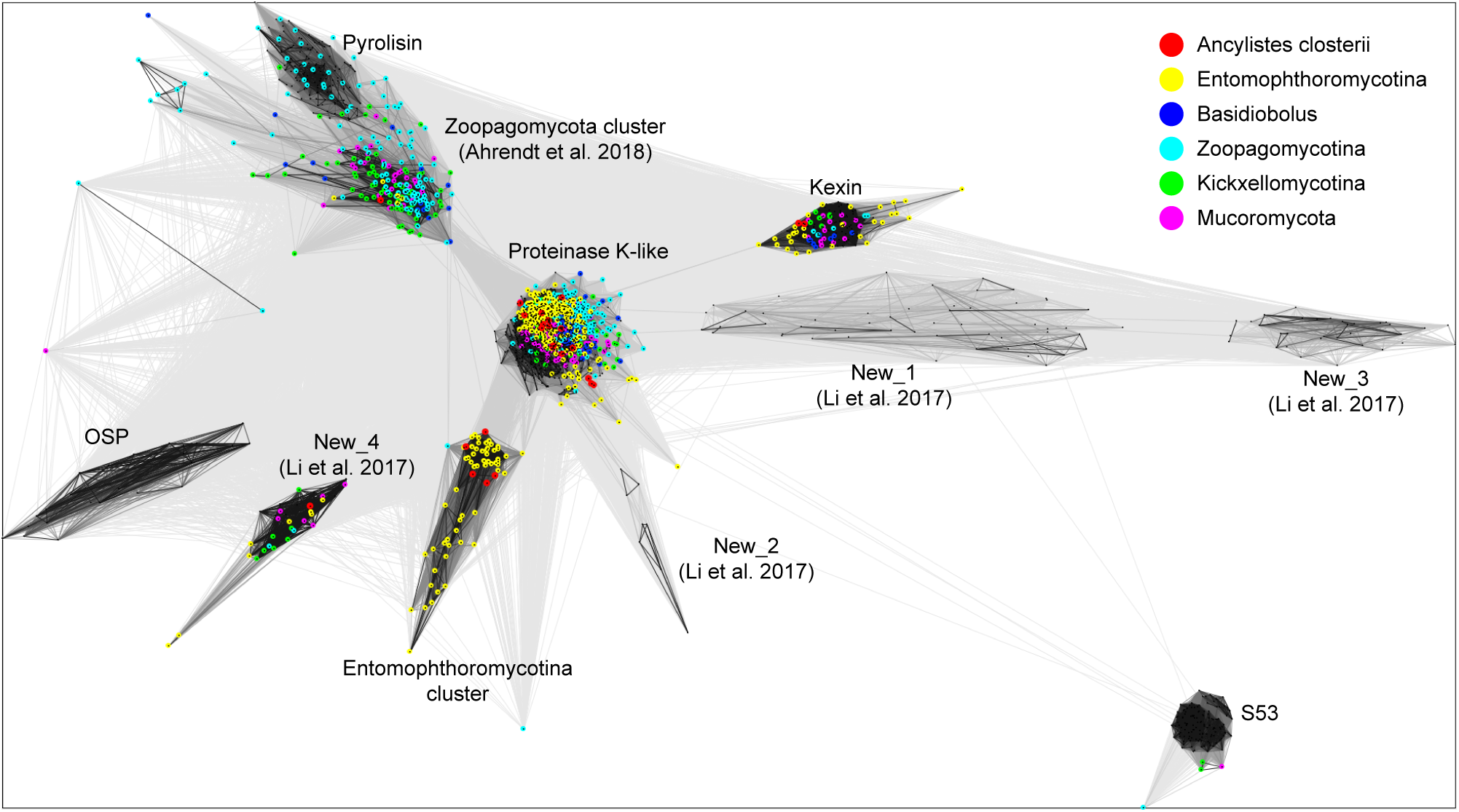
Clustering of fungal subtilase amino acid sequences. Names of groups are according to Li et al. (2017) and Ahrendt et al. (2018). The sequences from zygomycete genomes are indicated by colored circles.

Hybrid histidine kinases (HHKs) are major sensing proteins in Fungi, which are composed of three main regions: the variable N-terminal sensing domain, the central regions of histidine kinase A (HisKA) and histidine kinase-like ATPase catalytic (HATPase_c) domains, and the C-terminal receiver domain (Defosse et al. 2015). In *A. closterii*, four HHKs were found (Fig. 7A). A phylogenetic analysis showed that *A. closterii* HHKs were categorized into two clades (Fig. S12): one is in the Fungal HHK Group III (Defosse et al. 2015) and the others in an Entomophthoromycotina-specific clade. The Group III HHKs were characterized by multiple HAMP domains (Fig. 7A) and found in all zygomycete subphyla examined (Fig. S12, Table S5). The Entomophthoromycotina-specific HHKs were characterized by 1 or 2 PAS domains but some genes lacked it (Fig. 7A, Table S5). Some early diverging fungi possess ethylene receptor homologs, composed of core HHK domains and three transmembrane regions involved in ethylene reception (Hérivaux et al. 2017). A homology search of the ethylene receptor region showed that all Entomophthoromycotina fungi except for *E*. *muscae* possessed putative ethylene receptor homologs (Fig. 7A, Table S5). All detected genes had transmembrane regions with residues essential for ethylene reception (Wang et al. 2007) and a single GAF domain (Fig. 7A, B). The receiver domain was lacking in all Entomophthoromycotina genes (Fig. 7A). The histidine kinase region was absent in two *A. closterii* genes and incomplete in other Entomophthoromycotina (Fig. 7A).

**Fig. 7.**
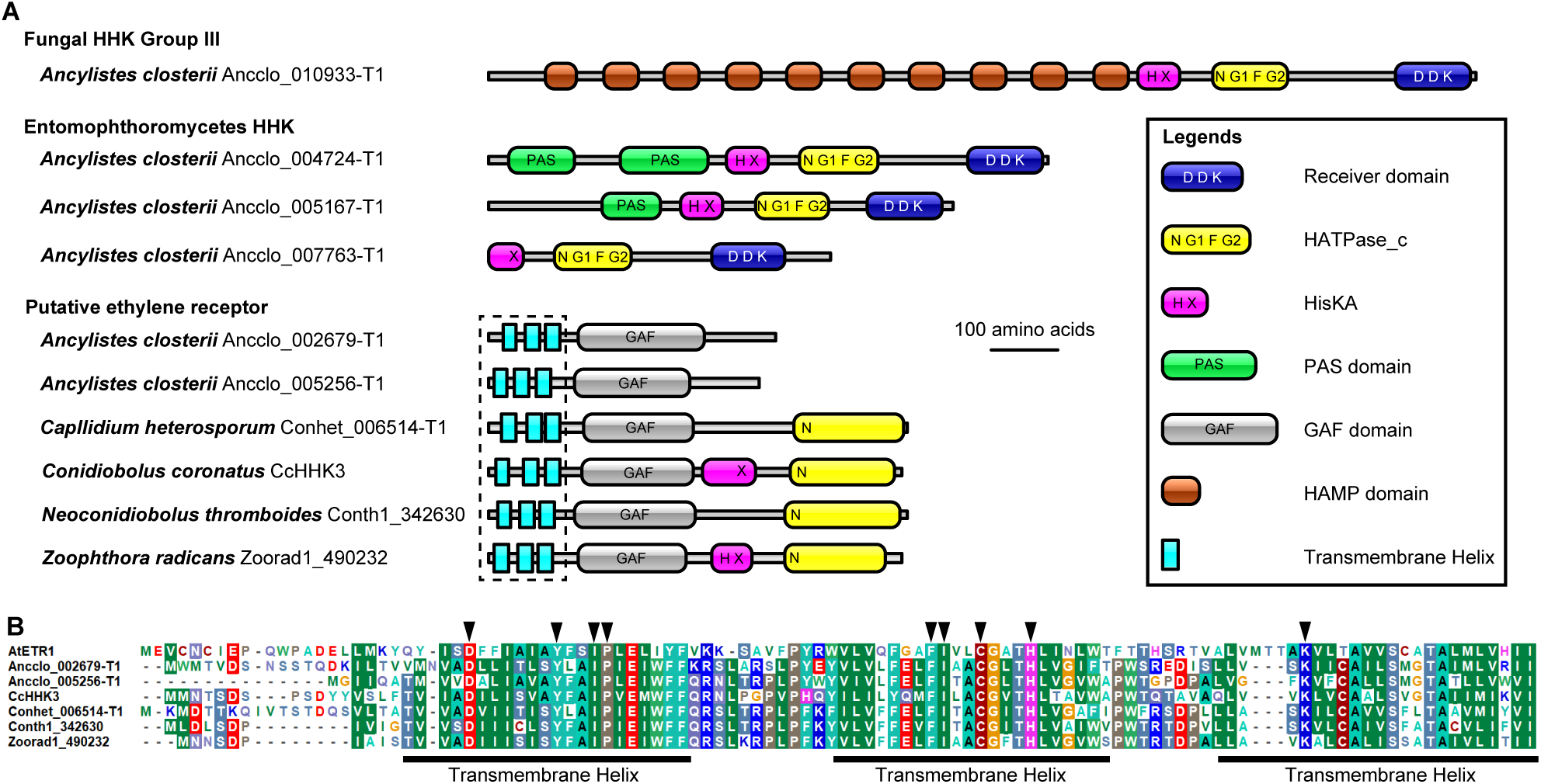
Hybrid histidine kinases (HHKs) and putative ethylene receptors (ETRs) found in Entomophthoromycotina genomes. (*A*) Domain architectures of four HHKs in *Ancylistes closterii* and ETRs found in *A. closterii* and other Entomophthoromycotina. (*B*) Alignment of amino acid sequences of three transmembrane regions of ETRs. AtETR1 is an ETR of *Arabidopsis thaliana*. Arrowheads indicate residues essential for ethylene reception shown in Wang et al. (2007).

Translation elongation factor 1α (EF-1α) is an essential protein involved in the translation process (Negrutskii & El’skaya 1998). Although EF-1α is a highly conserved protein in eukaryotes, it is replaced with a related protein with the same function, elongation factor-like (EFL) protein, in some lineages (Keeling & Inagaki 2004). In Fungi, some early diverging fungi including Entomophthoromycotina possess EFL (Kamikawa et al. 2013; Amses et al. 2022). A homology search showed that *A. closterii* possessed EFL but not EF-1α. In zygomycetes, EFL was found in Entomophthoromycotina taxa as well as *Calcarisporiella thermophila* (Mucoromycotina) and *Basidiobolus* spp., which possessed both EF-1α and EFL (Fig. 8). All other zygomycetes had EF-1α.

**Fig. 8.**
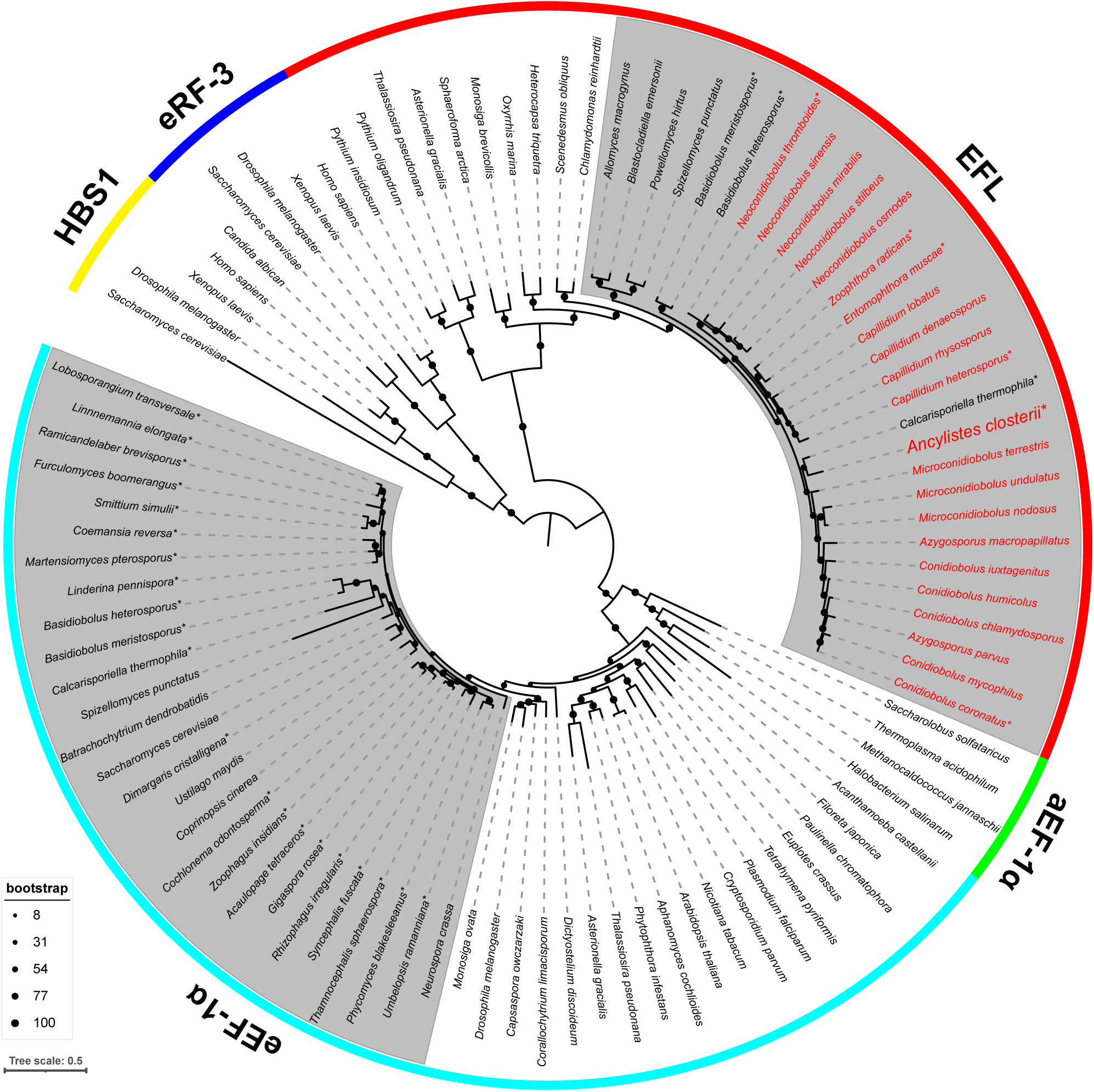
Maximum likelihood tree of eukaryotic elongation factor 1α (eEF-1α), elongation factor-like genes (EFL), and their related genes (aEF-1α = archaeal EF-1α, eRF-3 = eukaryotic release factor 3, HBS1 = heat shock protein 70 subfamily B suppressor 1). Fungal clades of eEF-1α and EFL are indicated by a gray background color. Sequences from zygomycete fungi are indicated by asterisks on each taxa name. Sequences from Entomophthoromycotina taxa are indicated by a red font label.

## Discussion

### Position of *A. closterii* and phylogeny of Entomophthoromycotina

Our phylogenetic analyses of the 18S-28S rDNA and genome-scale datasets showed *A. closterii* actually belongs to Entomophthoromycotina despite its great ecological distinctiveness. Our phylogenetic analyses also strongly supported the recent taxonomic revisions of Ancylistaceae (Nie et al. 2020; Gryganskyi et al. 2022). *Ancylistes closterii* was sister to the clade of Entomophthoraceae + paraphyletic *Conidiobolus* s.l. (Fig. 3), and alternative topologies such as the monophyly of Ancylistaceae as well as *Conidiobolus* s.l. were rejected in the AU test (Table S3). Another lineage, Batkoaceae including the genus *Batkoa* and some *Conidiobolus*-like taxa (Gryganskyi et al. 2022), was positioned outside of *A. closterii* in our 18S-28S rDNA phylogeny (Fig. S3). However, two previous analyses showed different positions, sister to Entomophthoraceae (Gryganskyi et al. 2022) or sister to Entomophthoraceae + Capillidiaceae + Neoconidiobolaceae (Möckel et al. 2022). These incongruences were probably caused by abnormally divergent rDNA sequences of Batkoaceae. Genome sequences of Batkoaceae are required to infer robust phylogeny of this family.

We considered that increased sampling of early-diverging Entomophthoromycotina might help stabilize the phylogenetic position of *Basidiobolus*. The phylogenetic position of *Basidiobolus* has long been controversial despite large genome-scale phylogenies. The recent phylogenetic studies showed the same position as our analysis, the sister position to all other Zoopagomycota lineages (Galindo et al. 2021; Strassert & Monaghan 2022; Wang et al. 2023). However, some studies reported alternative positions such as sister to Entomophthoromycotina (Spatafora et al. 2016; Wang et al. 2020) and Mucoromycota (Li et al. 2021). Our analyses revealed that the latter position of *Basidiobolus* could be caused by fast-evolving sites in the alignment and/or an overly simple model of amino acid substitution. Actually, a simple site-homogeneous model (LG+G4) was used in the concatenated alignment analysis by Li et al. (2021). Our analyses showed that *Basidiobolus* is at the earliest branch of Zoopagomycota and the alternative placements inside the phylum were rejected by the AU test. However, one of the two *Basidiobolus* genomes (*B. heterosporus*) is highly fragmented. The phylogeny of *Basidiobolus* needs to be examined carefully with more high-quality genomes as well as robust taxon sampling.

### Unique and shared genomic traits of *A. closterii* among Zoopagomycota lineages

*Ancylistes* is enigmatic as a member of Zoopagomycota because it is the only algal parasite in the phylum, which is composed of taxa primarily associated with animals and other fungi (Spatafora et al. 2016). Our comparative analyses of zygomycete fungi revealed that *Ancylistes* possessed unique enzymes putatively important for parasitism on algae. However, it also shares lineage-specific traits with other Entomophthoromycotina.

PCWDEs are involved in hydrolyzing polymers contained in plant cell walls, such as cellulose, hemicellulose, and pectin, and some of them have important roles in pathogenicity in plant pathogenic fungi (Zhao et al. 2014; Kubicek et al. 2014). Our investigation showed that three gene families/subfamilies, GH44, GH45, and PL3_2 were uniquely enriched in *Ancylistes*. In these, GH44 was unique for *Ancylistes* among zygomycete fungi although *Ancylistes* has only one copy of this gene. The family GH44 formerly included only bacterial endoglucanases and xyloglucanases (Warner et al. 2011) and was recently found in Fungi but restricted to the phylum Basidiomycota (Sun et al. 2022). Our surveys showed that some zoosporic fungal lineages also possess this xyloglucanase family. All fungal GH44 genes were clustered into a monophyletic clade in the gene tree (Fig. S9), indicating the ancestral acquisition of this gene family in Fungi but further surveys are required. Xyloglucan, a substrate of GH44 xyloglucanase, is a kind of hemicellulose and one of the major components of plant cell walls (Pauly et al. 2013). The charophyte green algae including *Closterium* (Zygnematophyceae), which are related to land plants, also have xyloglucan in their cell wall (Mikkelsen et al. 2021). Therefore, the GH44 gene in *Ancylistes* could be important for invasion into the host *Closterium* cell.

Subtilases are the most abundant and diverse serine proteases in Fungi (Muszewska et al. 2017). Recently, two groups of subtilases specific to Zoopagomycota lineages were found (Ahrendt et al. 2018; Arnesen et al. 2018). One of them is the “Zoopagomycota cluster” represented by genes from mycoparasitic lineages in Zoopagomycotina and Kickxellomycotina (Ahrendt et al. 2018). Our clustering analysis showed that this group of subtilases was shared by Mucoromycota lineages as well as Zoopagomycota including *Ancylistes*. Another group is specific to Entomophthoromycotina with the one exception of *Zoophagus insidians* (Zoopagomycotina) in our analysis. This unique group was formerly reported as a new group of subtilases restricted to bacteria, oomycetes, Entomophthoromycotina fungi, and two fungi of the Rozellomycota + Microsporidia lineage (Arnesen et al. 2018). Although the function of this unique subtilase is currently unknown, it is widely shared by Entomophthoromycotina taxa, from algal parasites to insect pathogens.

HHKs in Fungi are involved in the reception and signal transduction of various environmental stimuli and are essential for the physiological processes of fungal cells (Hérivaux et al. 2016). Fungal HHKs are classified into 16 groups (Defosse et al. 2015). *Ancylistes closterii* possessed one gene of the fungal group III HHK which was also found in most other zygomycete fungi. The fungal group III HHK is the most abundant HHK in Fungi and characterized by multiple HAMP domains in the N-terminal region (Defosse et al. 2015). The well-known function of the group III HHK is osmosensing but it is also essential for morphogenesis and virulence of pathogenic fungi (Defosse et al. 2015; Hérivaux et al. 2016). We also found a unique HHK group restricted to Entomophthoromycotina taxa including *Ancylistes*. HHKs of this group are characterized by one or two PAS domains in the N-terminal region but some of them lack this domain. In our gene tree, this Entomophthoromycotina-specific HHK group was sister to the fungal group IX which has three PAS domains (Defosse et al. 2015). The function of the group IX HHK has been examined in the rice blast fungus *Magnaporthe oryzae* (Jacob et al. 2014). They showed that deletion of the corresponding gene did not affect fungal growth under culture conditions but reduced virulence to the host. The Entomophthoromycotina-specific HHKs could be important for parasitism on algae and insects. However, it should be noted that saprotrophic taxa, *Conidiobolus* s.l., also possess this group of HHK. Another group of HHK found in *Ancylistes* was the putative ethylene receptor, which is characterized by three transmembrane regions for ethylene binding (Wang et al. 2007). Hérivaux et al. (2017) showed that some early diverging fungi have phytohormone receptors such as ethylene and cytokinin receptors. Our surveys revealed that all Entomophthoromycotina taxa except for *E. muscae* possessed ethylene receptor (ETR) homologs. As with the gene of *C. coronatus* (Hérivaux et al. 2017), these ETRs in Entomophthoromycotina had a GAF domain but their histidine kinase region was incomplete, and the receiver domain was lacking. In particular, two genes of *Ancylistes* lacked all histidine kinase regions. We also found that two Zoopagomycotina taxa, *Acaulopage tetraceros* and *Cochlonema odontosperma* have ETRs with complete histidine kinase regions (Table S4). Although the phylogeny of Entomophthoromycotina ETRs could not be examined due to their incomplete histidine kinase regions, all fungal ETRs could share an ancestor, and independent losses and/or gene reductions occurred (Hérivaux et al. 2017). Ethylene is known to be involved in the interaction between arbuscular mycorrhizal fungi and plants (Torres de Los Santos et al. 2011; Foo et al. 2016). Ju et al. (2015) showed that charophyte algae possess an ethylene hormone system similar to that of land plants. Therefore, the ETRs in *Ancylistes* could be important for interaction with the host *Closterium* belonging to the charophyte green algae. As with Entomophthoromycotina ETRs, some of the plant ETRs have incomplete histidine kinase regions (Gallie 2015). In rice, the ethylene receptor OsERS2, via its GAF domain, inhibits the kinase activity of MHZ1/OsHK1 which is essential for ethylene response signaling (Zhao et al. 2020). The ETRs found in *Ancylistes* possibly have similar functions with their GAF domain.

The translation elongation factor 1α (EF-1α) gene is essential for the translation process and is often used in phylogenetic analyses of fungal lineages (Tanabe et al. 2004; James et al. 2006). However, it is known that EF-1α is replaced with a related gene with the same function, elongation factor-like (EFL) gene in some fungal lineages (Kamikawa et al. 2013). Our survey strongly suggests that all Entomophthoromycotina taxa including *Ancylistes* have EFL instead of EF-1α. The gene sequences annotated as “elongation factor 1α” were used in some recent taxonomic and phylogenetic studies of *Conidiobolus* s.l. (Nie et al. 2020; Cai et al. 2021; Nie et al. 2021). Our gene tree analysis showed that these elongation factor genes from *Conidiobolus* s.l. are actually EFL.

## Conclusion

In the present study, we successfully established a dual culture of *A. closterii* and its host, which enabled us to characterize this enigmatic fungus in detail. Our phylogenetic analyses showed that *Ancylistes* is sister to *Conidiobolus* s.l. and insect pathogenic Entomophthoraceae. These results support the recent revision of Ancylistaceae, in which *Conidiobolus* s.l. is excluded from the family. However, the phylogenetic position of Batkoaceae could not be determined with strong support and will require genome data of Batkoaceae. Investigations of further poorly studied Entomophthoromycotina taxa such as Completoriaceae and Meristacraceae (Humber 2012) should also be pivotal to resolving the phylogeny of the group. Especially, the former family solely represented by *Completoria complens* is important because it is a parasite of fern prothallia (Tucker 1981).

*Ancylistes* is an abnormal fungus because it is the only algal parasite in Zoopagomycota represented by mainly animal-associated taxa. However, our comparative analyses revealed that *Ancylistes* shares many traits with other Entomophthoromycotina taxa such as possession of EFL instead of EF-1α and expansions of lineage-specific subtilases and HHKs. *Ancylistes* could have evolved as an algal parasite from a saprotrophic or animal-associated Entomophthoromycotina fungus-like ancestor. This may have been facilitated by specific retention of ancestral fungal unique PCWDEs and ethylene receptors that are important for parasitism and interaction with algae.

## Materials and Methods

### Isolation, Culturing, and Microscopic Observation

*Ancylistes closterii* parasitic on a green alga *Closterium* sp. was found in a water sample collected at a small pond in Ann Arbor, MI, USA (42°15’10.5”N, 83°42’00.0”W) on 24 July 2019. Because *A. closterii* is considered an obligate parasite, it was cultivated as a dual culture of fungus and its host. First, the host alga, *Closterium* sp., was isolated from the sample collected at another pond (42°16’58.3”N, 83°47’35.5”W) on 14 July 2019. A single algal cell was isolated under a Nikon TMS Inverted microscope (Nikon, Tokyo, Japan) using a manually prepared drawn-out glass capillary pipette. The isolated cell was transferred into a 24-well plate containing WC medium (Guillard & Lorenzen 1972) and brought into a culture (strain KSA123). Subsequently, an algal cell infected by fungal hyphae was isolated and transferred into a 24-well plate containing the host strain KSA123. The fungus-alga dual culture (strain KS117) was maintained by transferring the infected culture to fresh host culture at 7–14 days intervals. All strains were incubated at 20 °C and under LED lighting.

The morphology of *A. closterii* stain KS117 was observed and photographed using Zeiss Axio Imager A2 (differential interference contrast or epifluorescence) equipped with Zeiss AxioCam MRc camera. Resting spore formation was induced by incubating the culture under dark conditions. Conidial formation was induced by pipetting 1–2 mL of fungus-alga dual culture onto an 1% agar plate of WC medium. To observe nuclei in the cells of *A. closterii*, cells were stained with SYBR™ Green I (ThermoFisher) (0.01% in final concentration).

### DNA Extraction and Whole Genome Sequencing

Because strain KS117 contains non-target organisms (host alga and contaminated bacteria), a suspension of purified fungal conidia was prepared for DNA sequencing. Infected algae (2 mL) were inoculated onto 20 plates of WC medium (1% agar) and each plate was sealed with Parafilm M laboratory film (Bemis). After 3 days of incubation at 20 °C and under LED lighting, forcibly discharged conidia of *A. closterii* sticking to the lids of plates were observed. Conidia were collected by suspending them in 0.1% Tween 20, centrifuging at 1,000×g for 5 min, and removing the supernatant. Genomic DNA was extracted by 2× CTAB extraction protocol (James et al. 2008). A genomic library was prepared using the NEBNext Ultra DNA Library Prep Kit (New England BioLabs) and sequenced on an Illumina NovaSeq (S4 Flow Cell) at the U. Michigan Advanced Genomics Core. We obtained 640,219,350 paired-end reads (2 × 151 bp).

### Genome Assembly and Gene Prediction

Read quality was assessed using FastQC v0.11.5 (https://www.bioinformatics.babraham.ac.uk/projects/fastqc/). Illumina adapter sequences and low-quality sequences were trimmed using Trimmomatic v0.36 (Bolger et al. 2014) with the following options: ILLUMINACLIP:Nebnext.fa:3:30:10 LEADING:20 TRAILING:20 SLIDINGWINDOW:4:20 MINLEN:30. Trimmed paired-end reads were assembled using Platanus assembler v1.2.4 (Kajitani et al. 2014) with default options. Scaffolds smaller than 1,000 bp were excluded. The completeness of the assembly was assessed using BUSCO v4 (Manni et al. 2021) with the fungi_odb10 database. Gene prediction was performed using the Funannotate v1.7.4 pipeline (https://github.com/nextgenusfs/funannotate). Assembly was soft-masked by “funannotate mask” with default options. Gene models were predicted by the “funannotate predict” command, which uses Genemark-ES (Lomsadze et al. 2005), AUGUSTUS (Stanke & Waack 2003), GlimmerHMM (Majoros et al. 2004), and SNAP (Korf 2004) for gene prediction and EVidenceModeler (Haas et al. 2008) for integration of prediction results.

### Phylogenetic analysis of 18S and 28S rDNA

A rDNA operon sequence of *A. closterii* was assembled from the trimmed paired-end reads using GetOrganelle (Jin et al. 2020) with the “fungus_nr” database. From the obtained rDNA contig, 18S and 28S sequences were extracted using Barrnap (https://github.com/tseemann/barrnap). A dataset of Entomophthoromycotina and other major fungal phyla/subphyla was prepared (Table S1). Sequences were aligned using MAFFT v7.487 (Katoh & Standley 2013) with the “L-INS-i” method, and the alignment was trimmed using trimAl v1.2 (Capella-Gutiérrez et al. 2009) with the “gappyout” method. A maximum likelihood (ML) tree was inferred with IQ-TREE 2 (Minh et al. 2020). The best model of each alignment was examined using ModelFinder (Kalyaanamoorthy et al. 2017) implemented in the IQ-TREE 2. According to the corrected Akaike information criterion (AICc), GTR+F+R6 and GTR+F+R5 models were selected for 18S and 28S, respectively. An ML analysis was run with a partitioned model (Chernomor et al. 2016). The branch supports were assessed with standard nonparametric bootstrap analysis (100 replicates). The tree was visualized with FigTree (https://github.com/rambaut/figtree).

### Phylogenomic analyses

For phylogenomic analyses, proteome data of 49 fungal genomes were used (Table S2). Two Rozellomycota taxa (*Paramicrosporidium saccamoebae* and *Rozella allomycis*) were selected as the outgroup. Because proteome data were not available for *A*. *tetraceros*, *C*. *odontosperma*, *Z*. *insidians* (Zoopagomycotina), and *C*. *heterosporum* (Entomophthoromycotina), gene predictions were done using Funannotate v.1.7.4. The “protein mode” of BUSCO version 4 (Manni et al. 2021) was run for each proteome using the fungi_odb10 database. Single copy BUSCO genes shared by 80% of 49 taxa were retrieved, aligned (muscle v3) (Edgar 2004), trimmed (trimAl), and integrated into a concatenated alignment using version 4 of the “BUSCO_phylogenomics” pipeline (https://github.com/jamiemcg/BUSCO_phylogenomics, (McGowan et al. 2020)). The supermatrix alignment consisted of 185,578 amino acids (aa) of 452 genes. Preliminary ML analyses were performed using IQ-TREE 2 with site-homogeneous (LG+F+R10) and site-heterogeneous (LG+C20+F+G+PMSF (Wang et al. 2018) using the guide tree inferred with LG+F+R10 model) models. A fast-site removal assay was performed using the “fast_site_remover.py” command of PhyloFisher (Tice et al. 2021). ML tree was inferred for each new alignment using IQ-TREE 2 with LG+F+R10 model, and ultrafast bootstrap values (Hoang et al. 2018) of relevant clades were recorded (Fig. S6). Subsequent phylogenetic analyses were performed on the alignment from which the 50,000 fastest evolving sites were removed. An ML tree was inferred using IQ-TREE 2 with LG+C60+F+G+PMSF model (a tree inferred with LG+F+R10 model was used as a guide tree) and 100 replicates of standard nonparametric bootstrap analysis were run. We also conducted a Bayesian analysis with the CAT-Poisson+G4 model using PhyloBayes MPI v1.8 (Lartillot et al. 2013). Four Markov Monte Carlo chains were run for at least 7,500 generations. The four chains were moderately converged after the removal of the first 3,750 generations (effective size ≥ 57 and discrepancies ≤ 0.259882, calculated by the “tracecomp” command), and a consensus tree was obtained. To evaluate alternative tree topologies (Table S3), an approximately unbiased (AU) test (Shimodaira 2002) was performed using IQ-TREE 2 with LG+C60+F+G model according to its “Advanced tutorial (http://www.iqtree.org/doc/Advanced-Tutorial#tree-topology-tests)”.

### Investigation of specific genes in *A. closterii* and other zygomycete fungi

To examine the general differences between *A. closterii* and other Entomophthoromycotina taxa, GO enrichment analysis was performed. We also focused on specific gene families relevant to the transition between algal and arthropod hosts. We investigated PCWDEs, subtilases, HHKs, and EF1α/EFL genes in the selected 27 zygomycete genomes. To infer gene trees, alignment, trimming, and ML analysis were performed using MAFFT v7.487, trimAl v1.2, and IQ-TREE 2, respectively. Trees were visualized and edited with iTOL (Letunic & Bork 2021). To obtain GO term annotations and to investigate subtilases and HHKs, annotations inferred with InterProScan v5.55-88.0 (Jones et al. 2014) were used.

We performed GO enrichment analysis using the BiNGO plugin (Maere et al. 2005) in Cytoscape (Shannon et al. 2003). GO term annotations were retrieved from the InterProScan results of *A. closterii*, *C. heterosporum*, *C. coronatus*, *N. thromboides*, *E. muscae*, and *Z. radicans*. The genes without GO term annotations were excluded. A hypergeometric test with the Benjamini & Hochberg False Discovery Rate correction (significant level of 0.01) was performed.

To investigate PCWDEs, carbohydrate-active enzymes (CAZymes) in each zygomycete proteome were searched using the HMMER search option in the dbCAN2 meta server (Zhang et al. 2018). PCWDEs repertoire of zygomycete fungi was shown as a heatmap using the “pheatmap” package (Kolde 2019) in R (R Core Team 2022). Gene trees were inferred for GH44, GH45, and PL3_2 (Table S6-S8).

For subtilases, genes annotated with the Pfam domain PF00082 (Subtilase family) were retrieved. Subfamily classifications of zygomycete subtilases were visualized using CLANS (Frickey & Lupas 2004) which shows clustering of proteins based on pairwise similarity. The zygomycete subtilase sequences were added to the dataset of Li et al. (2017) and an all-against-all BLAST search was performed using the CLANS web-utility in the MPI Bioinformatics Toolkit (Gabler et al. 2020).

HHKs in each zygomycete proteome were identified based on annotations with three functional domains: Histidine Kinase A domain (PF00512), ATPase catalytic (HATPase_c) domain (PF02518), and Response regulator receiver domain (PF00072). Ethylene receptor homologs were searched by BLASTP searches using some ethylene receptor gene sequences found by Hérivaux et al. (2017). Domain structures of HHKs in *A. closterii* and ethylene receptor genes in Entomophthoromycotina fungi were visualized using IBS 2.0 (Xie et al. 2022). Phylogenetic relationships of zygomycete HHKs were inferred using the modified dataset of Hérivaux et al. (2017).

The EF1α and EFL genes were searched by BLASTP (Altschul et al. 1990) against each zygomycete proteome using the amino acid sequences of EF1α from *Coemansia reversa* (Accession no. PIA15833.1) and EFL from *Conidiobolus coronatus* (Accession no. KXN73350.1) as queries. Identification of the detected genes was verified by phylogenetic analysis. Initially, the sequences except for the short sequences of *Conidiobolus* s.l., which were deposited as “elongation factor 1-alpha” (Nie et al. 2020; Cai et al. 2021) (Table S9), were aligned. The aforementioned short sequences were added into the trimmed alignment using the “--add” and “--keeplength” options of MAFFT v7.487.

## Acknowledgements

K.S. was supported by JSPS Overseas Research Fellowship (No. 201960485) and JSPS KAKENHI Grant Number JP22KJ1394. This project was funded, in part, by the U.S. National Science Foundation grant DEB-1441677. T.Y.J. is a fellow of CIFAR program Fungal Kingdom: Threats & Opportunities. We thank Adrian Tsang for allowing us to use the genome of *Calcarisporiella thermophila*.

## Data Availability

The raw sequence data, genome assembly, and annotations of *Ancylistes closterii* have been deposited in the DNA Data Bank of Japan: BioProject (PRJDB17127), Illumina sequence reads (DRR516680), assembly and annotations (BTXW01000000). Additional data generated in this study such as gene sequences, alignments, and phylogenetic trees are available in a figshare repository (Project No. 189093).

