## Supplementay_Information for "Genomic analysis of *Ancylistes closterii*, an enigmatic alga parasitic fungus in the arthropod-associated Entomophthoromycotina"

**Supplementary Table Legends**

Table S1. List of taxa and sequences used for the phylogenetic analysis of 18S-28S rDNA.

Table S2. List of genomic data used for phylogenomic analysis.

Table S3. Results of approximately unbiased test.

Table S4. List of over- and underrepresented gene ontology terms in *Ancylistes closterii* compared with other Entomophthoromycotina taxa.

Table S5. List of hybrid histidine kinases and ethylene receptor homologs investigated in this study.

Table S5. List of hybrid histidine kinases and ethylene receptor homologs investigated in this study.

Table S6. List of GH44 sequences used for phylogenetic analysis.

Table S7. List of GH45 sequences used for phylogenetic analysis.

Table S8. List of PL3_2 sequences used for phylogenetic analysis.

Table S9. List of sequences of translation elongation factor 1-alpha, elongation factor-like, and their related genes used for phylogenetic analysis.

**Supplementary Figures and Legends**

Figure S1. Nuclei of *Ancylistes closterii* stained with SYBR green. Images *A*, *E*, *G*, *I*, and *K* were observed with differential interference contrast microscopy. Images *B–D*, *F*, *H*, *J*, and *L* were observed with fluorescence microscopy. (*A–D*) Multinucleated conidium. Observation of the same conidium at different focuses shows up to 14 nuclei. (*E*, *F*) Developing external hyphae including nine nuclei. (*G*, *H*) Early developmental stage of internal hyphae with 12 nuclei in the host cell. The large ellipsoidal stained region is the host nuclei. (*I*, *J*) Developing multinucleated internal hyphae in the host cell. (*K*, *L*) Segmented internal hyphae in the host cell. Image *L* indicates four or five nuclei are positioned in one segment. Bars = 10 µm.

Figure S2. Ploidy estimation by kmer and allele frequency analyses using short read sequences and genome assembly of *Ancylistes closterii*.

Figure S3. Maximum likelihood (ML) tree reconstructed with a concatenated dataset of 18S and 28S rDNA. Values next to nodes indicate ML bootstrap values. Nodes supported 100% bootstrap value were indicated by black circles. Double slashes on branches indicate that length is reduced by half.

Figure S4. Preliminary maximum likelihood (ML) tree reconstructed with a concatenated dataset (185,578 amino acids of 452 genes). The tree was inferred with the LG+F+R10 model using IQ-TREE2, and support values of nodes were calculated with ultrafast bootstrapping analysis (1,000 replicates). Only bootstrap values less than 100 were indicated next to nodes.

Figure S5. Preliminary maximum likelihood (ML) tree reconstructed with a concatenated dataset (185,578 amino acids of 452 genes). The tree was inferred with the LG+C20+F+G+PMSF model using IQ-TREE2, and support values of nodes were calculated with ultrafast bootstrapping analysis (1,000 replicates). Only bootstrap values less than 100 were indicated next to nodes.

Figure S6. Result of fast-evolving site removal assay. The fastest-evolving sites were removed every 10,000 sites per step. The ultrafast bootstrap values for each interested grouping were recorded by the LG+F+R10 model analysis.

Figure S7. Bayesian tree reconstructed with a concatenated dataset after removal of the 50,000 fastest evolving sites (135,578 amino acids of 452 genes). The tree was inferred with the CAT-Poisson+G4 model using PhyloBayes MPI v1.8. Only Bayesian posterior probability less than 1 was indicated next to one node.

Figure S8. Hierarchical organizations of overrepresented (*A*) and underrepresented (*B*) gene ontology (GO) terms in *Ancylistes closterii* compared to other Entomophthoromycotina taxa. Colored circles indicate significantly overrepresented (yellow to orange) and underrepresented (light blue to dark blue) GO terms.

Figure S9. Maximum likelihood tree of glycoside hydrolase family 44 (GH44) genes from Fungi and bacteria. The tree was inferred with the Q.pfam+F+R7 model selected by the ModelFinder in the IQ-TREE2. Ultrafast bootstrap values are indicated by circles on the nodes.

Figure S10. Maximum likelihood tree of glycoside hydrolase family 45 (GH45) genes from Fungi, Metazoa, and bacteria. The tree was inferred with the Q.pfam+R6 model selected by the ModelFinder in the IQ-TREE2. Ultrafast bootstrap values are indicated by circles on the nodes.

Figure S11. Maximum likelihood tree of polysaccharide lyase subfamily 2 of family 3 (PL3_2) genes from Fungi, Metazoa, Oomycota, and bacteria. The tree was inferred with the WAG+I+G4 model selected by the ModelFinder in the IQ-TREE2. Ultrafast bootstrap values are indicated by circles on the nodes.

Figure S12. Maximum likelihood tree of hybrid histidine kinase genes from Fungi. The tree was inferred with the Q.pfam+F+R8 model selected by the ModelFinder in the IQ-TREE2. Ultrafast bootstrap values are indicated by circles on the nodes.

### Figure S1

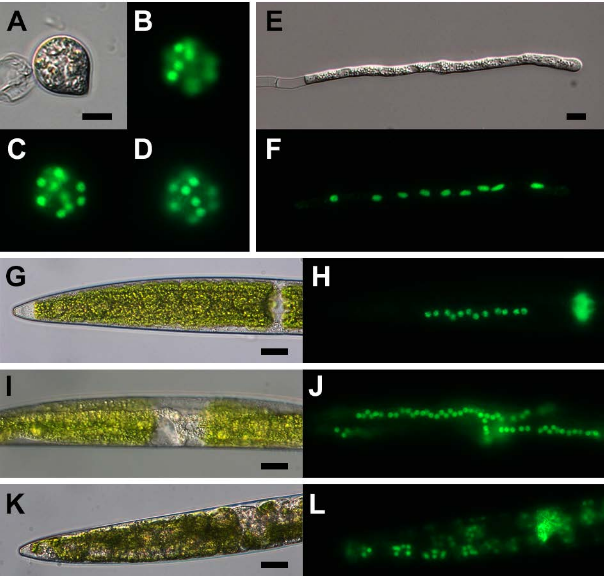

Figure S2

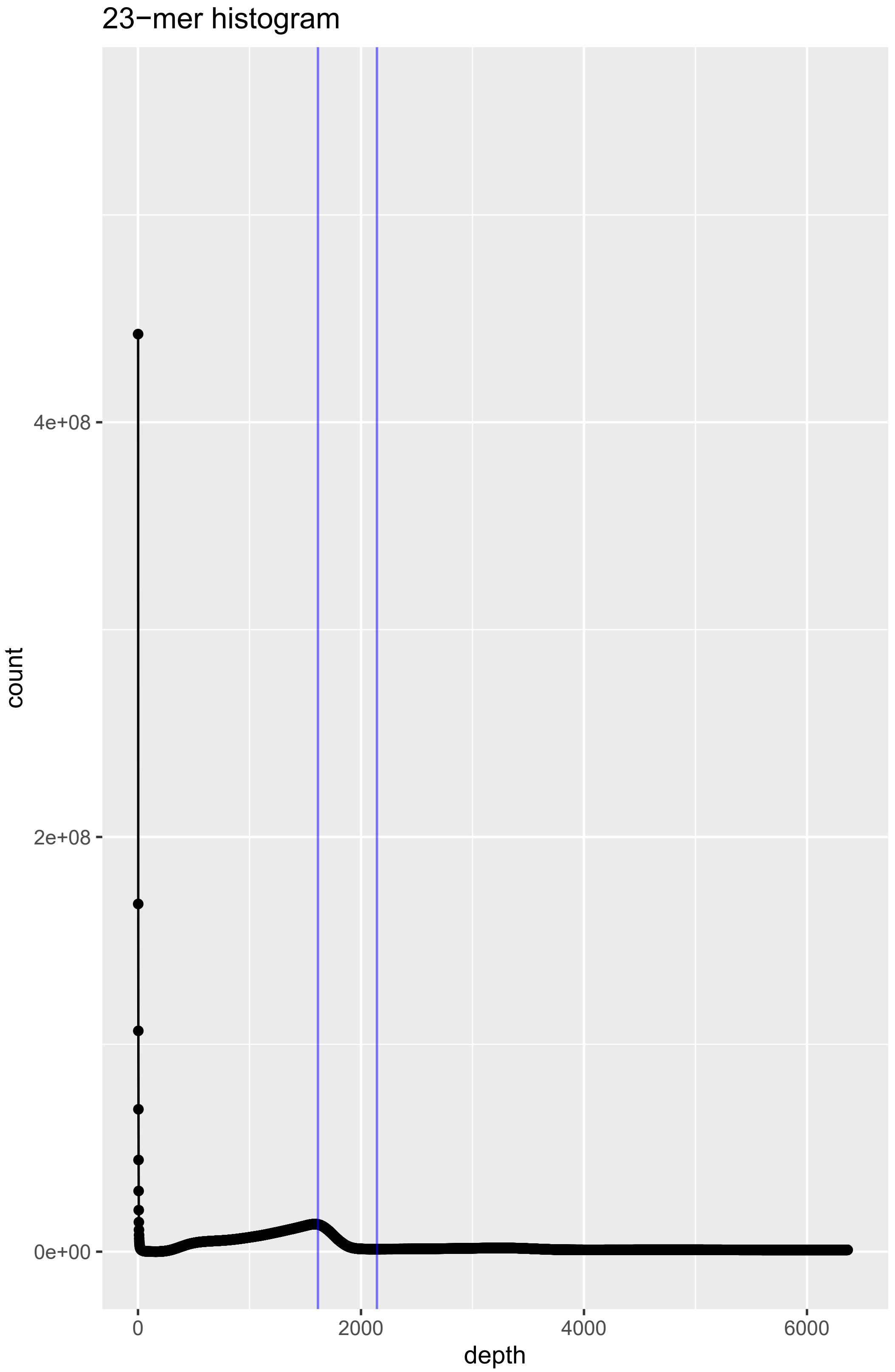
23−mer histogram
count
4e+08
2e+08
0e+00
0
2000
4000
6000
depth

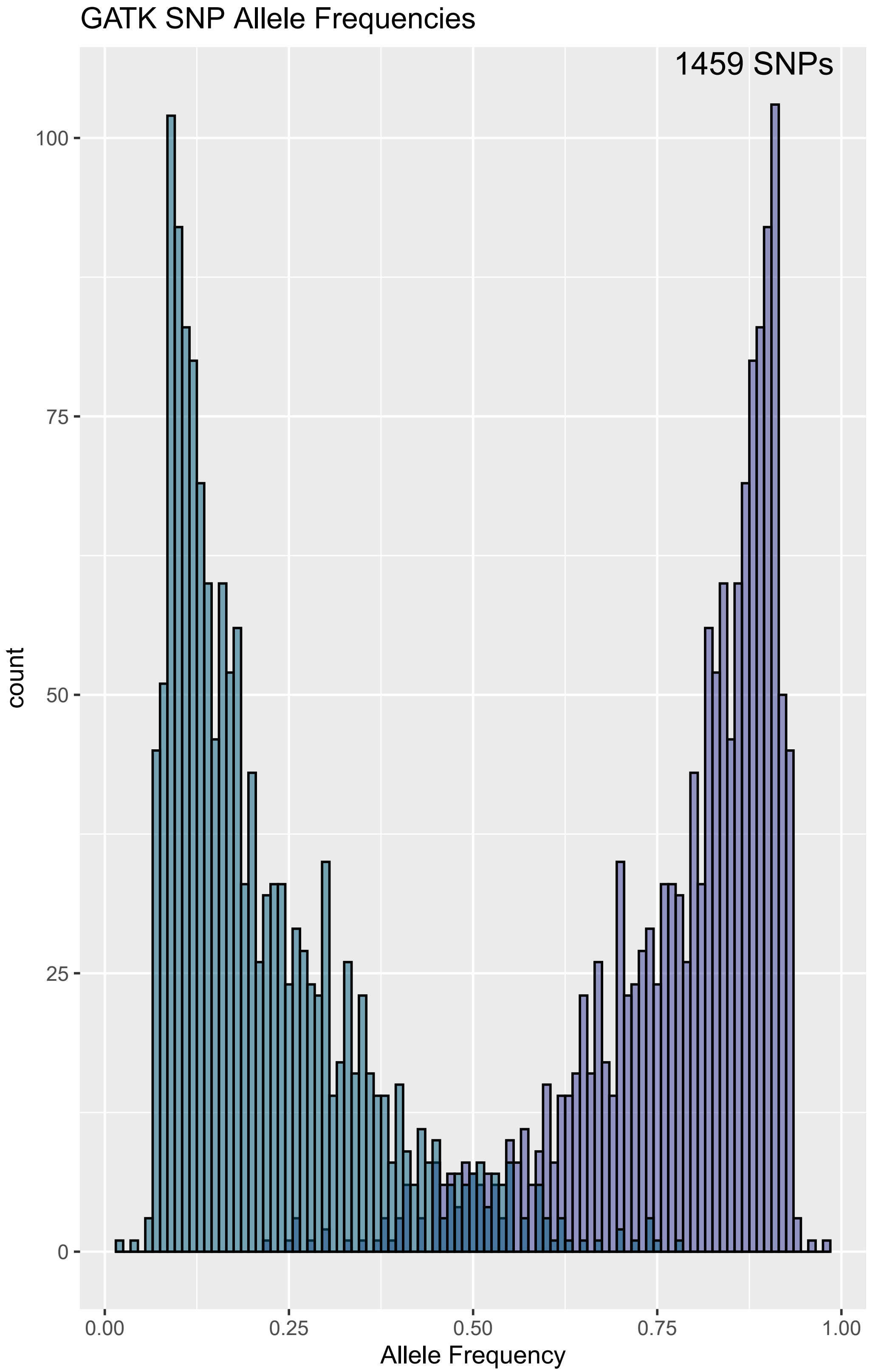
GATK SNP Allele Frequencies
1459 SNPs
count
100
75
50
25
0
0.00
0.25
0.50
0.75
1.00
Allele Frequency

### Figure S3

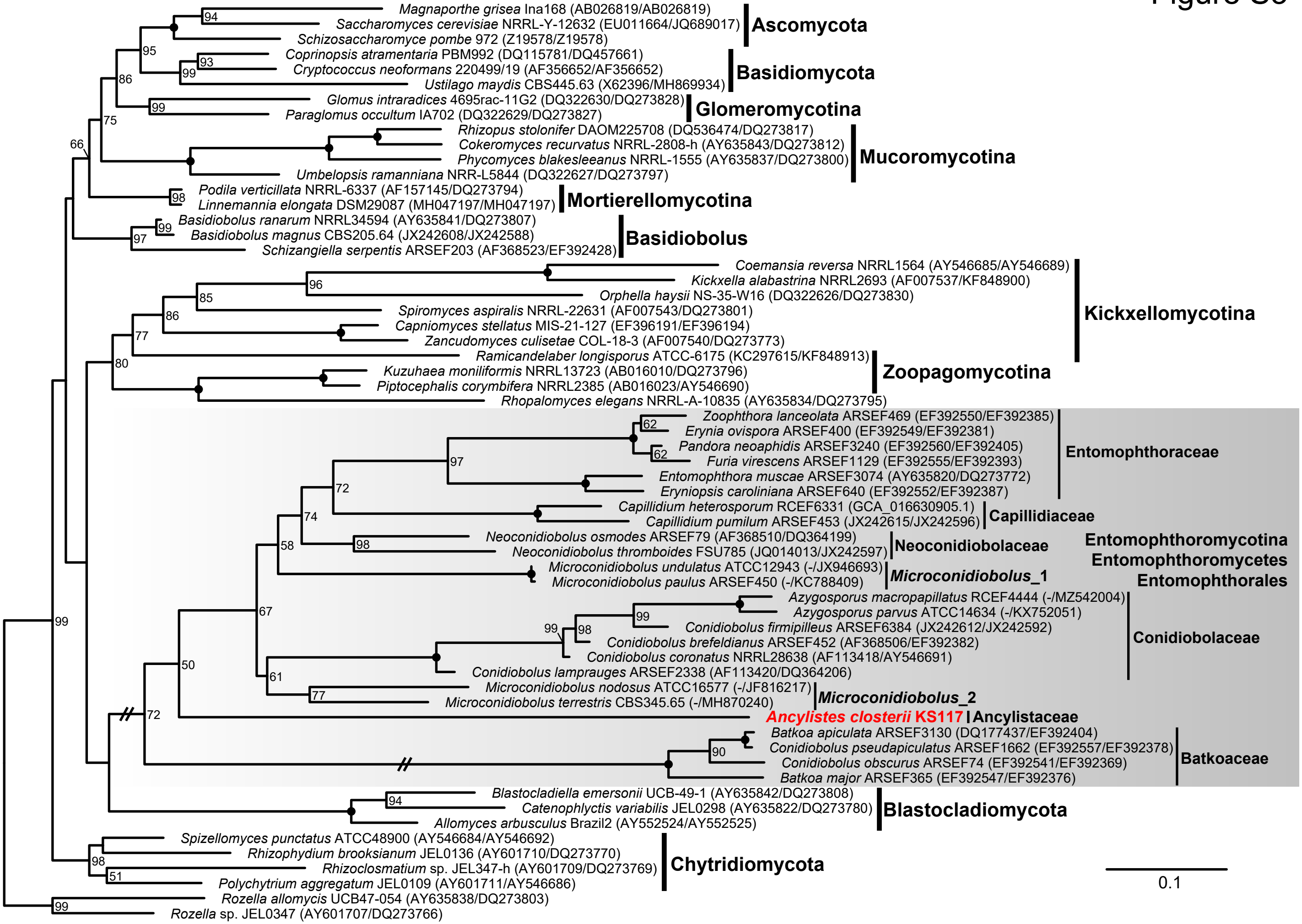

Magnaporthe grisea Ina168 (AB026819/AB026819)
Saccharomyces cerevisiae NRRL-Y-12632 (EU011664/JQ689017)
Schizosaccharomyce pombe 972 (Z19578/Z19578)
Ascomycota
Coprinopsis atramentaria PBM992 (DQ115781/DQ457661)
Cryptococcus neoformans 220499/19 (AF356652/AF356652)
Ustilago maydis CBS445.63 (X62396/MH869934)
Basidiomycota
Glomus intraradices 4695rac-11G2 (DQ322630/DQ273828)
Paraglomus occultum IA702 (DQ322629/DQ273827)
Glomeromycotina
Rhizopus stolonifer DAOM225708 (DQ536474/DQ273817)
Cokeromyces recurvatus NRRL-2808-h (AY635843/DQ273812)
Phycomyces blakesleeanus NRRL-1555 (AY635837/DQ273800)
Umbelopsis ramanniana NRR-L5844 (DQ322627/DQ273797)
Mucoromycotina
Podila verticillata NRRL-6337 (AF157145/DQ273794)
Linnemannia elongata DSM29087 (MH047197/MH047197)
Mortierellomycotina
Basidiobolus ranarum NRRL34594 (AY635841/DQ273807)
Basidiobolus magnus CBS205.64 (JX242608/JX242588)
Schizangiella serpentis ARSEF203 (AF368523/EF392428)
Basidiobolus
Coemansia reversa NRRL1564 (AY546685/AY546689)
Kickxella alabastrina NRRL2693 (AF007537/KF848900)
Orphella haysii NS-35-W16 (DQ322626/DQ273830)
Spiromyces aspiralis NRRL-22631 (AF007543/DQ273801)
Capniomyces stellatus MIS-21-127 (EF396191/EF396194)
Zancudomyces culisetae COL-18-3 (AF007540/DQ273773)
Ramicandelaber longisporus ATCC-6175 (KC297615/KF848913)
Kickxellomycotina
Kuzuhaea moniliformis NRRL13723 (AB016010/DQ273796)
Piptocephalis corymbifera NRRL2385 (AB016023/AY546690)
Rhopalomyces elegans NRRL-A-10835 (AY635834/DQ273795)
Zoopagomycotina
Zoophthora lanceolata ARSEF469 (EF392550/EF392385)
Erynia ovispora ARSEF400 (EF392549/EF392381)
Pandora neoaphidis ARSEF3240 (EF392560/EF392405)
Furia virescens ARSEF1129 (EF392555/EF392393)
Entomophthora muscae ARSEF3074 (AY635820/DQ273772)
Eryniopsis caroliniana ARSEF640 (EF392552/EF392387)
Entomophthoraceae
Capillidium heterosporum RCEF6331 (GCA_016630905.1)
Capillidium pumilum ARSEF453 (JX242615/JX242596)
Capillidiaceae
Neoconidiobolus osmodes ARSEF79 (AF368510/DQ364199)
Neoconidiobolus thromboides FSU785 (JQ014013/JX242597)
Neoconidiobolaceae
Microconidiobolus undulatus ATCC12943 (-/JX946693)
Microconidiobolus paulus ARSEF450 (-/KC788409)
Microconidiobolus_1
Entomophthoromycotina
Entomophthoromycetes
Entomophthorales
Azygosporus macropapillatus RCEF4444 (-/MZ542004)
Azygosporus parvus ATCC14634 (-/KX752051)
Conidiobolus firmipilleus ARSEF6384 (JX242612/JX242592)
Conidiobolus brefeldianus ARSEF452 (AF368506/EF392382)
Conidiobolus coronatus NRRL28638 (AF113418/AY546691)
Conidiobolus lamprauges ARSEF2338 (AF113420/DQ364206)
Conidiobolaceae
Microconidiobolus nodosus ATCC16577 (-/JF816217)
Microconidiobolus terrestris CBS345.65 (-/MH870240)
Microconidiobolus_2
Ancylistes closterii KS117
Ancylistaceae
Batkoa apiculata ARSEF3130 (DQ177437/EF392404)
Conidiobolus pseudapiculatus ARSEF1662 (EF392557/EF392378)
Conidiobolus obscurus ARSEF74 (EF392541/EF392369)
Batkoa major ARSEF365 (EF392547/EF392376)
Batkoaceae
Blastocladiella emersonii UCB-49-1 (AY635842/DQ273808)
Catenophlyctis variabilis JEL0298 (AY635822/DQ273780)
Allomyces arbusculus Brazil2 (AY552524/AY552525)
Blastocladiomycota
Spizellomyces punctatus ATCC48900 (AY546684/AY546692)
Rhizophydium brooksianum JEL0136 (AY601710/DQ273770)
Rhizoclosmatium sp. JEL347-h (AY601709/DQ273769)
Polychytrium aggregatum JEL0109 (AY601711/AY546686)
Chytridiomycota
Rozella allomycis UCB47-054 (AY635838/DQ273803)
Rozella sp. JEL0347 (AY601707/DQ273766)
94
95
93
99
86
99
75
66
98
99
97
96
85
86
77
80
62
62
97
72
74
98
58
67
99
99
98
61
77
50
72
90
94
98
51
99
99
0.1

### Figure S4

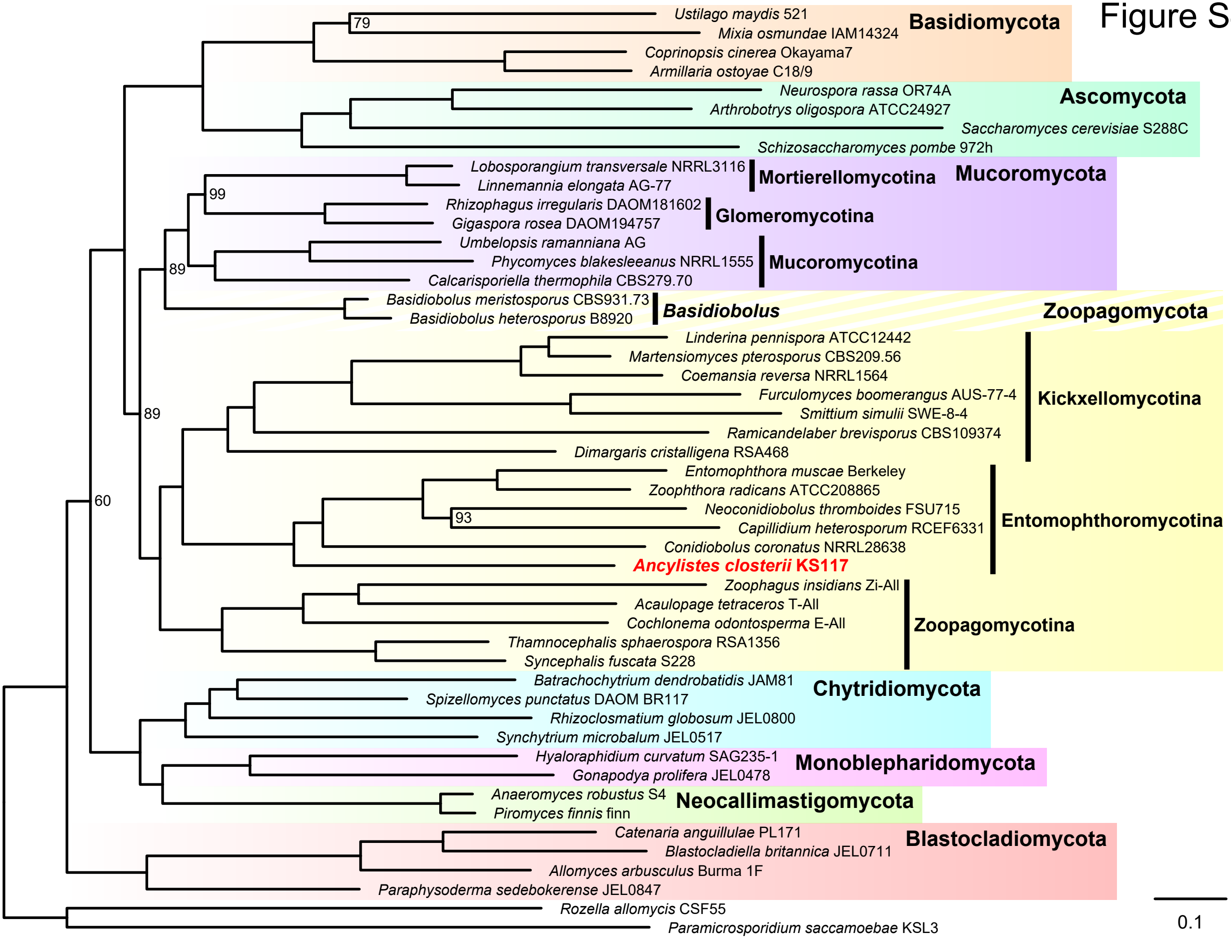

Basidiomycota
Ustilago maydis 521
Mixia osmundae IAM14324
79
Coprinopsis cinerea Okayama7
Armillaria ostoyae C18/9
Ascomycota
Neurospora rassa OR74A
Arthrobotrys oligospora ATCC24927
Saccharomyces cerevisiae S288C
Schizosaccharomyces pombe 972h
Mucoromycota
Mortierellomycotina
Lobosporangium transversale NRRL3116
Linnemannia elongata AG-77
99
Glomeromycotina
Rhizophagus irregularis DAOM181602
Gigaspora rosea DAOM194757
Mucoromycotina
Umbelopsis ramanniana AG
Phycomyces blakesleeanus NRRL1555
Calcarisporiella thermophila CBS279.70
89
Zoopagomycota
Basidiobolus
Basidiobolus meristosporus CBS931.73
Basidiobolus heterosporus B8920
Kickxellomycotina
Linderina pennispora ATCC12442
Martensiomyces pterosporus CBS209.56
Coemansia reversa NRRL1564
Furculomyces boomerangus AUS-77-4
Smittium simulii SWE-8-4
Ramicandelaber brevisporus CBS109374
Dimargaris cristalligena RSA468
89
60
Entomophthoromycotina
Entomophthora muscae Berkeley
Zoophthora radicans ATCC208865
Neoconidiobolus thromboides FSU715
93
Capillidium heterosporum RCEF6331
Conidiobolus coronatus NRRL28638
Ancylistes closterii KS117
Zoopagomycotina
Zoophagus insidians Zi-All
Acaulopage tetraceros T-All
Cochlonema odontosperma E-All
Thamnocephalis sphaerospora RSA1356
Syncephalis fuscata S228
Chytridiomycota
Batrachochytrium dendrobatidis JAM81
Spizellomyces punctatus DAOM BR117
Rhizoclosmatium globosum JEL0800
Synchytrium microbalum JEL0517
Monoblepharidomycota
Hyaloraphidium curvatum SAG235-1
Gonapodya prolifera JEL0478
Neocallimastigomycota
Anaeromyces robustus S4
Piromyces finnis finn
Blastocladiomycota
Catenaria anguillulae PL171
Blastocladiella britannica JEL0711
Allomyces arbusculus Burma 1F
Paraphysoderma sedebokerense JEL0847
Rozella allomycis CSF55
Paramicrosporidium saccamoebae KSL3
0.1

### Figure S5

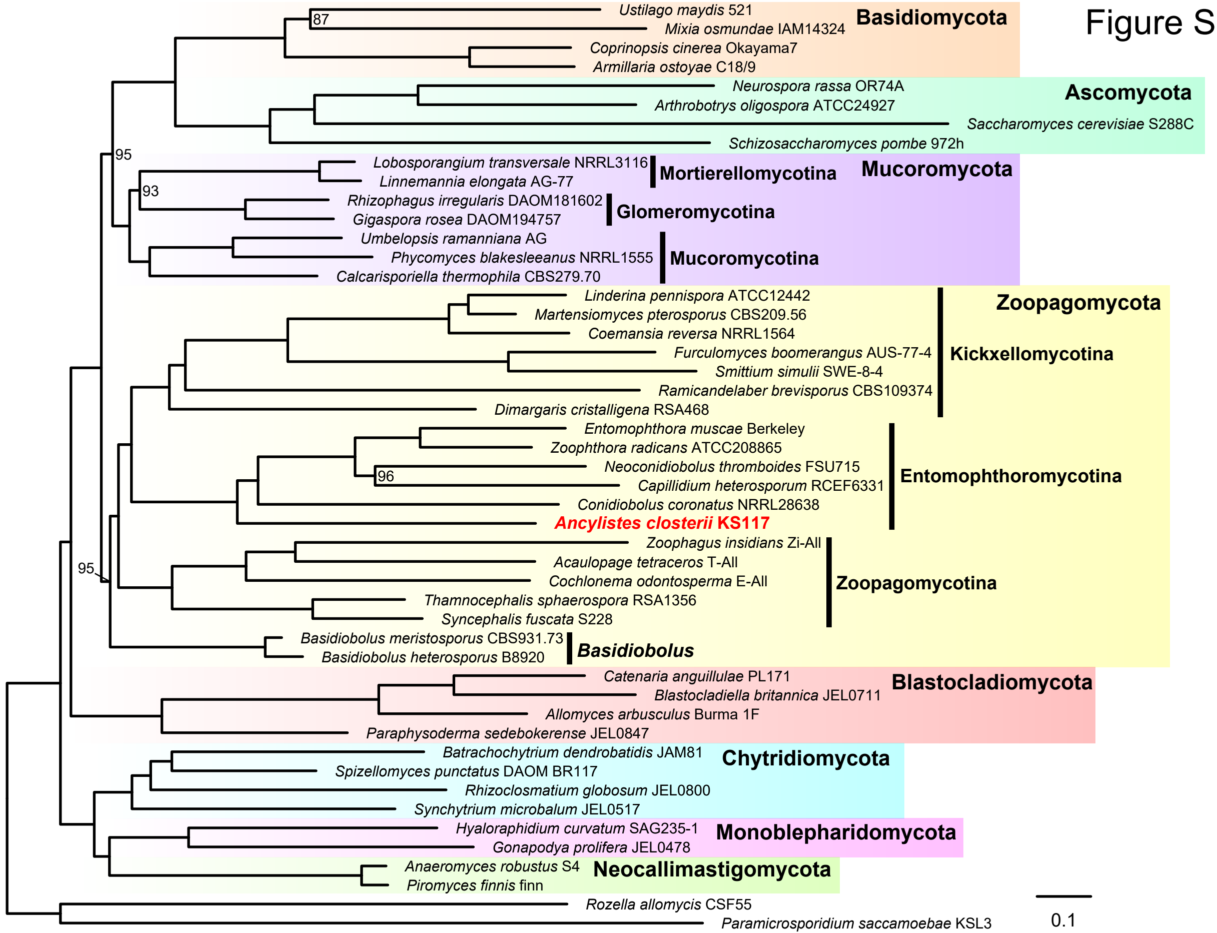
Basidiomycota
Ustilago maydis 521
Mixia osmundae IAM14324
Coprinopsis cinerea Okayama7
Armillaria ostoyae C18/9
87
Ascomycota
Neurospora rassa OR74A
Arthrobotrys oligospora ATCC24927
Saccharomyces cerevisiae S288C
Schizosaccharomyces pombe 972h
95
Mucoromycota
Lobosporangium transversale NRRL3116
Linnemannia elongata AG-77
Mortierellomycotina
93
Rhizophagus irregularis DAOM181602
Gigaspora rosea DAOM194757
Glomeromycotina
Umbelopsis ramanniana AG
Phycomyces blakesleeanus NRRL1555
Calcarisporiella thermophila CBS279.70
Mucoromycotina
Zoopagomycota
Linderina pennispora ATCC12442
Martensiomyces pterosporus CBS209.56
Coemansia reversa NRRL1564
Furculomyces boomerangus AUS-77-4
Smittium simulii SWE-8-4
Ramicandelaber brevisporus CBS109374
Dimargaris cristalligena RSA468
Kickxellomycotina
Entomophthora muscae Berkeley
Zoophthora radicans ATCC208865
Neoconidiobolus thromboides FSU715
96
Capillidium heterosporum RCEF6331
Conidiobolus coronatus NRRL28638
Ancylistes closterii KS117
Entomophthoromycotina
Zoophagus insidians Zi-All
Acaulopage tetraceros T-All
Cochlonema odontosperma E-All
Thamnocephalis sphaerospora RSA1356
Syncephalis fuscata S228
Zoopagomycotina
95
Basidiobolus meristosporus CBS931.73
Basidiobolus heterosporus B8920
Basidiobolus
Blastocladiomycota
Catenaria anguillulae PL171
Blastocladiella britannica JEL0711
Allomyces arbusculus Burma 1F
Paraphysoderma sedebokerense JEL0847
Chytridiomycota
Batrachochytrium dendrobatidis JAM81
Spizellomyces punctatus DAOM BR117
Rhizoclosmatium globosum JEL0800
Synchytrium microbalum JEL0517
Monoblepharidomycota
Hyaloraphidium curvatum SAG235-1
Gonapodya prolifera JEL0478
Neocallimastigomycota
Anaeromyces robustus S4
Piromyces finnis finn
Rozella allomycis CSF55
Paramicrosporidium saccamoebae KSL3
0.1

Figure S6

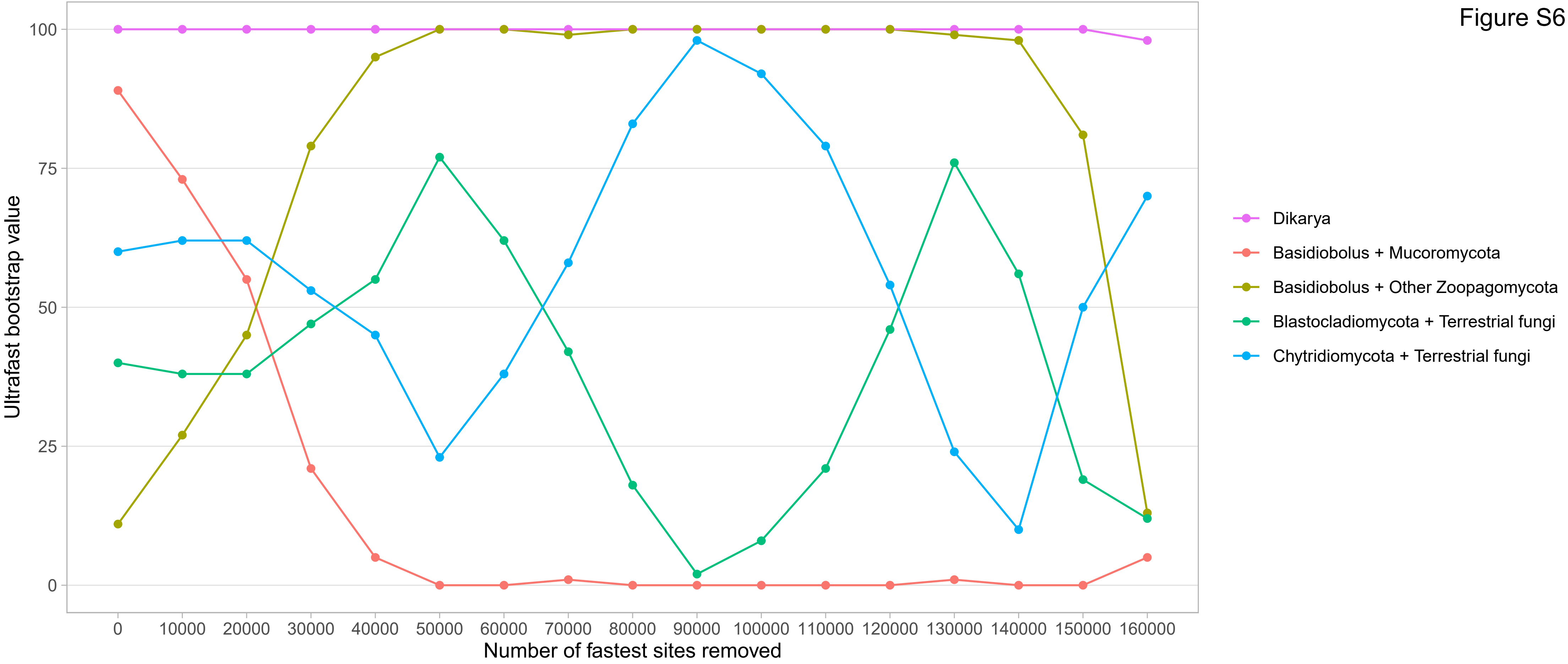

Ultrafast bootstrap value
100
75
50
25
0
0
10000
20000
30000
40000
50000
60000
70000
80000
90000
100000
110000
120000
130000
140000
150000
160000
Number of fastest sites removed
Dikarya
Basidiobolus + Mucoromycota
Basidiobolus + Other Zoopagomycota
Blastocladiomycota + Terrestrial fungi
Chytridiomycota + Terrestrial fungi

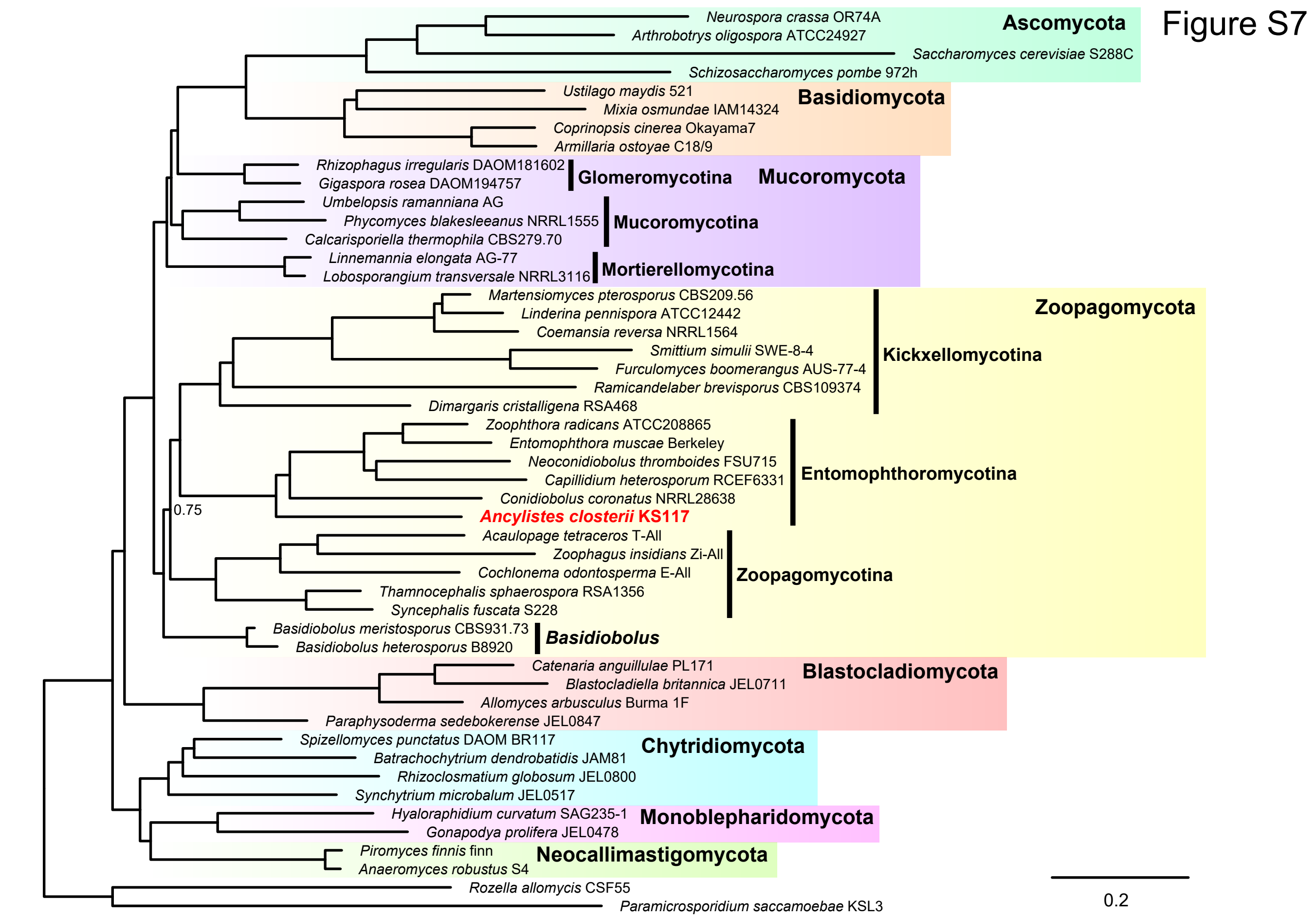

Figure S7
Ascomycota
Neurospora crassa OR74A
Arthrobotrys oligospora ATCC24927
Saccharomyces cerevisiae S288C
Schizosaccharomyces pombe 972h
Basidiomycota
Ustilago maydis 521
Mixia osmundae IAM14324
Coprinopsis cinerea Okayama7
Armillaria ostoyae C18/9
Mucoromycota
Rhizophagus irregularis DAOM181602
Gigaspora rosea DAOM194757
Glomeromycotina
Umbelopsis ramanniana AG
Phycomyces blakesleeanus NRRL1555
Calcarisporiella thermophila CBS279.70
Mucoromycotina
Linnemannia elongata AG-77
Lobosporangium transversale NRRL3116
Mortierellomycotina
Zoopagomycota
Martensiomyces pterosporus CBS209.56
Linderina pennispora ATCC12442
Coemansia reversa NRRL1564
Smittium simulii SWE-8-4
Furculomyces boomerangus AUS-77-4
Ramicandelaber brevisporus CBS109374
Dimargaris cristalligena RSA468
Kickxellomycotina
Zoophthora radicans ATCC208865
Entomophthora muscae Berkeley
Neoconidiobolus thromboides FSU715
Capillidium heterosporum RCEF6331
Conidiobolus coronatus NRRL28638
Ancylistes closterii KS117
Entomophthoromycotina
0.75
Acaulopage tetraceros T-All
Zoophagus insidians Zi-All
Cochlonema odontosperma E-All
Thamnocephalis sphaerospora RSA1356
Syncephalis fuscata S228
Zoopagomycotina
Basidiobolus meristosporus CBS931.73
Basidiobolus heterosporus B8920
Basidiobolus
Blastocladiomycota
Catenaria anguillulae PL171
Blastocladiella britannica JEL0711
Allomyces arbusculus Burma 1F
Paraphysoderma sedebokerense JEL0847
Chytridiomycota
Spizellomyces punctatus DAOM BR117
Batrachochytrium dendrobatidis JAM81
Rhizoclosmatium globosum JEL0800
Synchytrium microbalum JEL0517
Monoblepharidomycota
Hyaloraphidium curvatum SAG235-1
Gonapodya prolifera JEL0478
Neocallimastigomycota
Piromyces finnis finn
Anaeromyces robustus S4
Rozella allomycis CSF55
Paramicrosporidium saccamoebae KSL3
0.2

Figure S8

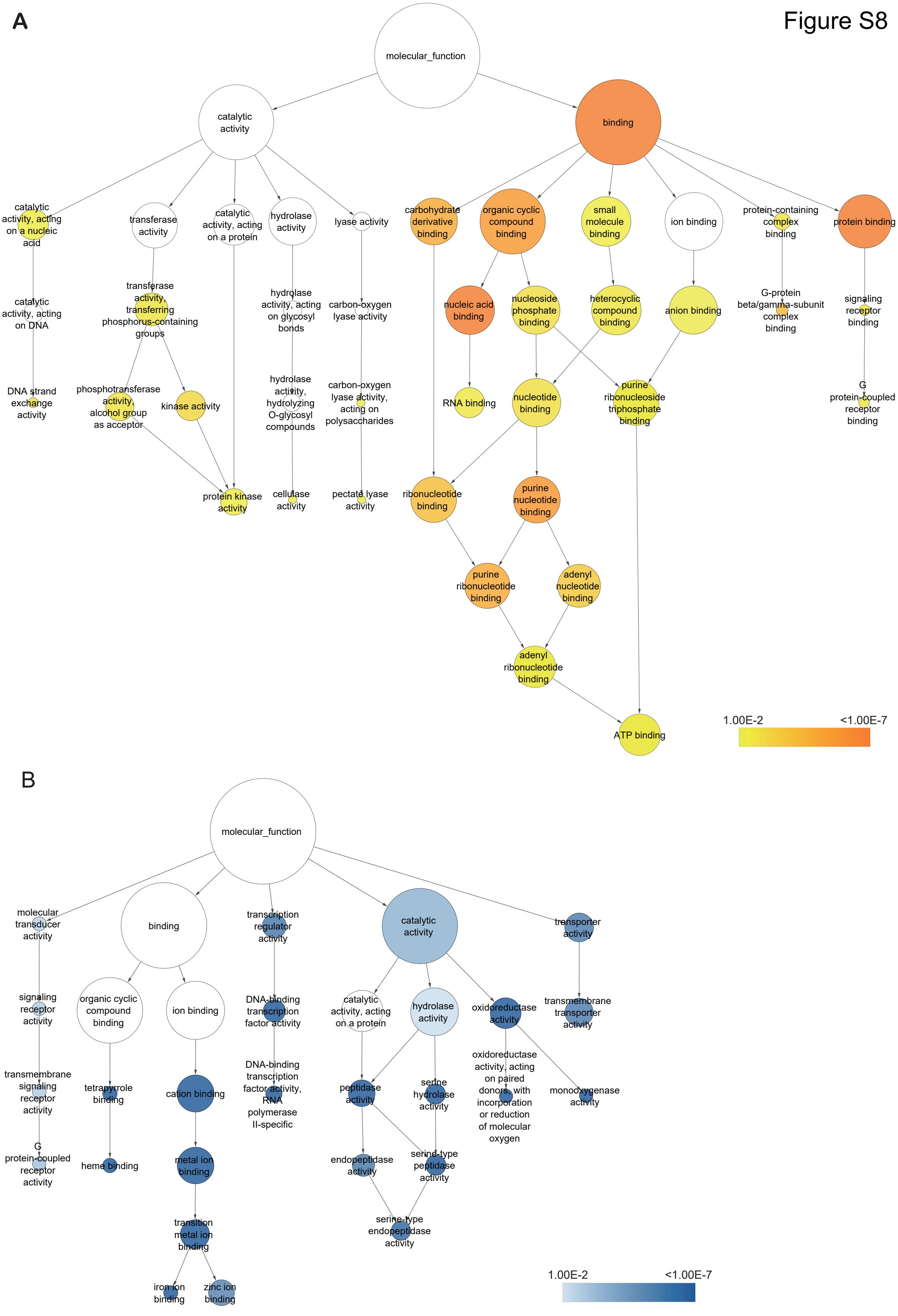

A
molecular_function
catalytic activity
binding
catalytic activity, acting on a nucleic acid
transferase activity
catalytic activity, acting on a protein
hydrolase activity
lyase activity
carbohydrate derivative binding
organic cyclic compound binding
small molecule binding
ion binding
protein-containing complex binding
protein binding
catalytic activity, acting on DNA
transferase activity, transferring phosphorus-containing groups
hydrolase activity, acting on glycosyl bonds
carbon-oxygen lyase activity
nucleic acid binding
nucleoside phosphate binding
heterocyclic compound binding
anion binding
G-protein beta/gamma-subunit complex binding
signaling receptor binding
DNA strand exchange activity
phosphotransferase activity, alcohol group as acceptor
kinase activity
hydrolase activity, hydrolyzing O-glycosyl compounds
carbon-oxygen lyase activity, acting on polysaccharides
RNA binding
nucleotide binding
purine ribonucleoside triphosphate binding
G protein-coupled receptor binding
protein kinase activity
cellulase activity
pectate lyase activity
ribonucleotide binding
purine nucleotide binding
purine ribonucleotide binding
adenyl nucleotide binding
adenyl ribonucleotide binding
ATP binding
1.00E-2
<1.00E-7
B
molecular_function
molecular transducer activity
binding
transcription regulator activity
catalytic activity
transporter activity
signaling receptor activity
organic cyclic compound binding
ion binding
DNA-binding transcription factor activity
catalytic activity, acting on a protein
hydrolase activity
oxidoreductase activity
transmembrane transporter activity
transmembrane signaling receptor activity
tetrapyrrole binding
cation binding
DNA-binding transcription factor activity, RNA polymerase II-specific
peptidase activity
serine hydrolase activity
oxidoreductase activity, acting on paired donors, with incorporation or reduction of molecular oxygen
monooxygenase activity
G protein-coupled receptor activity
heme binding
metal ion binding
endopeptidase activity
serine-type peptidase activity
serine-type endopeptidase activity
transition metal ion binding
iron ion binding
zinc ion binding
1.00E-2
<1.00E-7

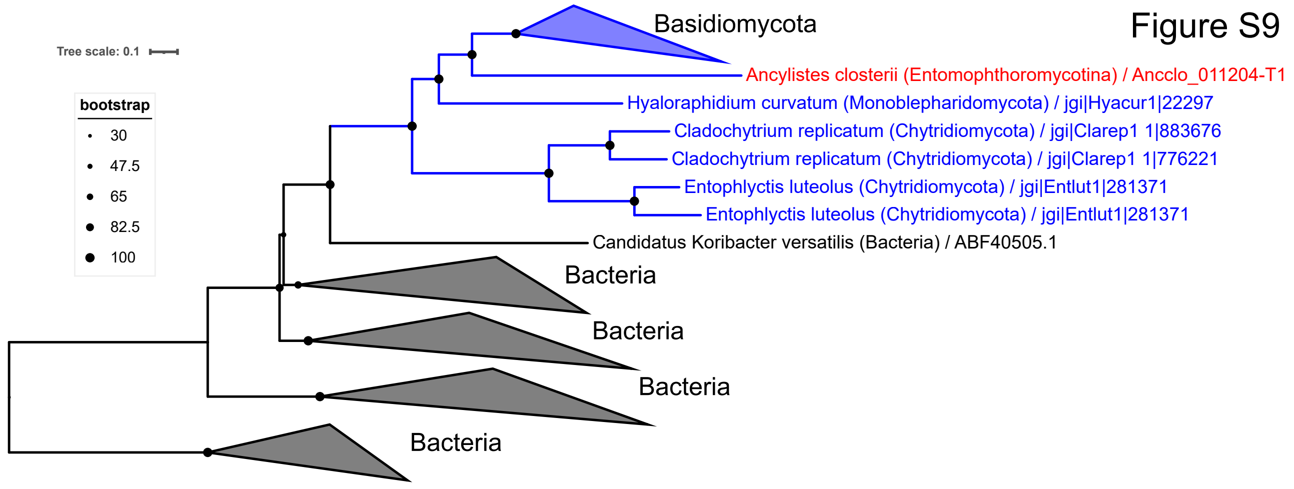
Figure S9
Tree scale: 0.1
bootstrap
30
47.5
65
82.5
100
Basidiomycota
Ancylistes closterii (Entomophthoromycotina) / Ancclo_011204-T1
Hyaloraphidium curvatum (Monoblepharidomycota) / jgi|Hyacur1|22297
Cladochytrium replicatum (Chytridiomycota) / jgi|Clarep1 1|883676
Cladochytrium replicatum (Chytridiomycota) / jgi|Clarep1 1|776221
Entophlyctis luteolus (Chytridiomycota) / jgi|Entlut1|281371
Entophlyctis luteolus (Chytridiomycota) / jgi|Entlut1|281371
Candidatus Koribacter versatilis (Bacteria) / ABF40505.1
Bacteria
Bacteria
Bacteria
Bacteria

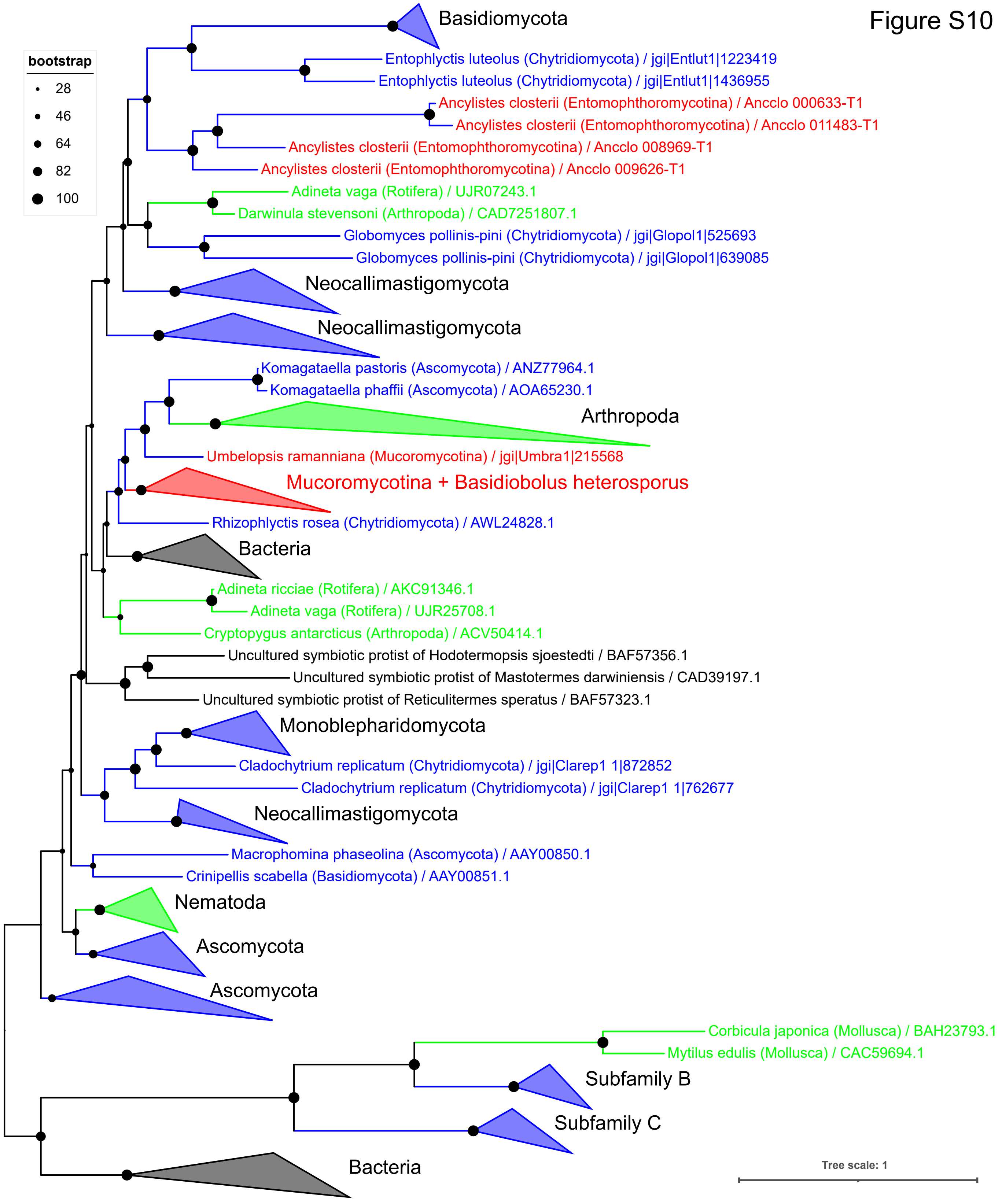
bootstrap
28
46
64
82
100
Basidiomycota
Entophlyctis luteolus (Chytridiomycota) / jgi|Entlut1|1223419
Entophlyctis luteolus (Chytridiomycota) / jgi|Entlut1|1436955
Ancylistes closterii (Entomophthoromycotina) / Ancclo 000633-T1
Ancylistes closterii (Entomophthoromycotina) / Ancclo 011483-T1
Ancylistes closterii (Entomophthoromycotina) / Ancclo 008969-T1
Ancylistes closterii (Entomophthoromycotina) / Ancclo 009626-T1
Adineta vaga (Rotifera) / UJR07243.1
Darwinula stevensoni (Arthropoda) / CAD7251807.1
Globomyces pollinis-pini (Chytridiomycota) / jgi|Glopol1|525693
Globomyces pollinis-pini (Chytridiomycota) / jgi|Glopol1|639085
Neocallimastigomycota
Neocallimastigomycota
Komagataella pastoris (Ascomycota) / ANZ77964.1
Komagataella phaffii (Ascomycota) / AOA65230.1
Arthropoda
Umbelopsis ramanniana (Mucoromycotina) / jgi|Umbra1|215568
Mucoromycotina + Basidiobolus heterosporus
Rhizophlyctis rosea (Chytridiomycota) / AWL24828.1
Bacteria
Adineta ricciae (Rotifera) / AKC91346.1
Adineta vaga (Rotifera) / UJR25708.1
Cryptopygus antarcticus (Arthropoda) / ACV50414.1
Uncultured symbiotic protist of Hodotermopsis sjoestedti / BAF57356.1
Uncultured symbiotic protist of Mastotermes darwiniensis / CAD39197.1
Uncultured symbiotic protist of Reticulitermes speratus / BAF57323.1
Monoblepharidomycota
Cladochytrium replicatum (Chytridiomycota) / jgi|Clarep1 1|872852
Cladochytrium replicatum (Chytridiomycota) / jgi|Clarep1 1|762677
Neocallimastigomycota
Macrophomina phaseolina (Ascomycota) / AAY00850.1
Crinipellis scabella (Basidiomycota) / AAY00851.1
Nematoda
Ascomycota
Ascomycota
Corbicula japonica (Mollusca) / BAH23793.1
Mytilus edulis (Mollusca) / CAC59694.1
Subfamily B
Subfamily C
Bacteria
Tree scale: 1

Figure S10

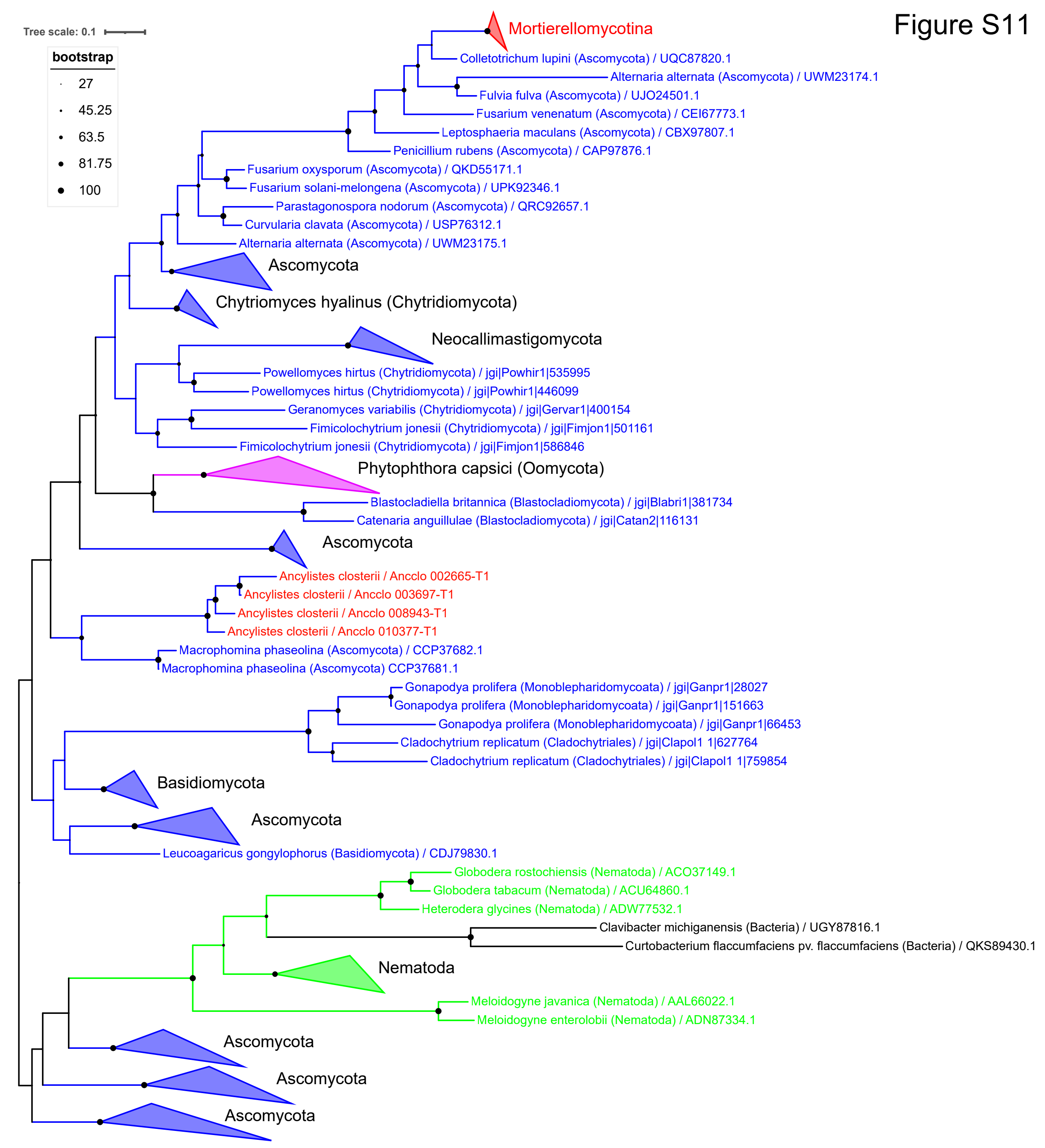

Figure S11
Tree scale: 0.1
bootstrap
27
45.25
63.5
81.75
100
Mortierellomycotina
Colletotrichum lupini (Ascomycota) / UQC87820.1
Alternaria alternata (Ascomycota) / UWM23174.1
Fulvia fulva (Ascomycota) / UJO24501.1
Fusarium venenatum (Ascomycota) / CEI67773.1
Leptosphaeria maculans (Ascomycota) / CBX97807.1
Penicillium rubens (Ascomycota) / CAP97876.1
Fusarium oxysporum (Ascomycota) / QKD55171.1
Fusarium solani-melongena (Ascomycota) / UPK92346.1
Parastagonospora nodorum (Ascomycota) / QRC92657.1
Curvularia clavata (Ascomycota) / USP76312.1
Alternaria alternata (Ascomycota) / UWM23175.1
Ascomycota
Chytriomyces hyalinus (Chytridiomycota)
Neocallimastigomycota
Powellomyces hirtus (Chytridiomycota) / jgi|Powhir1|535995
Powellomyces hirtus (Chytridiomycota) / jgi|Powhir1|446099
Geranomyces variabilis (Chytridiomycota) / jgi|Gervar1|400154
Fimicolochytrium jonesii (Chytridiomycota) / jgi|Fimjon1|501161
Fimicolochytrium jonesii (Chytridiomycota) / jgi|Fimjon1|586846
Phytophthora capsici (Oomycota)
Blastocladiella britannica (Blastocladiomycota) / jgi|Blabri1|381734
Catenaria anguillulae (Blastocladiomycota) / jgi|Catan2|116131
Ascomycota
Ancylistes closterii / Ancclo 002665-T1
Ancylistes closterii / Ancclo 003697-T1
Ancylistes closterii / Ancclo 008943-T1
Ancylistes closterii / Ancclo 010377-T1
Macrophomina phaseolina (Ascomycota) / CCP37682.1
Macrophomina phaseolina (Ascomycota) CCP37681.1
Gonapodya prolifera (Monoblepharidomycoata) / jgi|Ganpr1|28027
Gonapodya prolifera (Monoblepharidomycoata) / jgi|Ganpr1|151663
Gonapodya prolifera (Monoblepharidomycoata) / jgi|Ganpr1|66453
Cladochytrium replicatum (Cladochytriales) / jgi|Clapol1 1|627764
Cladochytrium replicatum (Cladochytriales) / jgi|Clapol1 1|759854
Basidiomycota
Ascomycota
Leucoagaricus gongylophorus (Basidiomycota) / CDJ79830.1
Globodera rostochiensis (Nematoda) / ACO37149.1
Globodera tabacum (Nematoda) / ACU64860.1
Heterodera glycines (Nematoda) / ADW77532.1
Clavibacter michiganensis (Bacteria) / UGY87816.1
Curtobacterium flaccumfaciens pv. flaccumfaciens (Bacteria) / QKS89430.1
Nematoda
Meloidogyne javanica (Nematoda) / AAL66022.1
Meloidogyne enterolobii (Nematoda) / ADN87334.1
Ascomycota
Ascomycota
Ascomycota

Figure S12

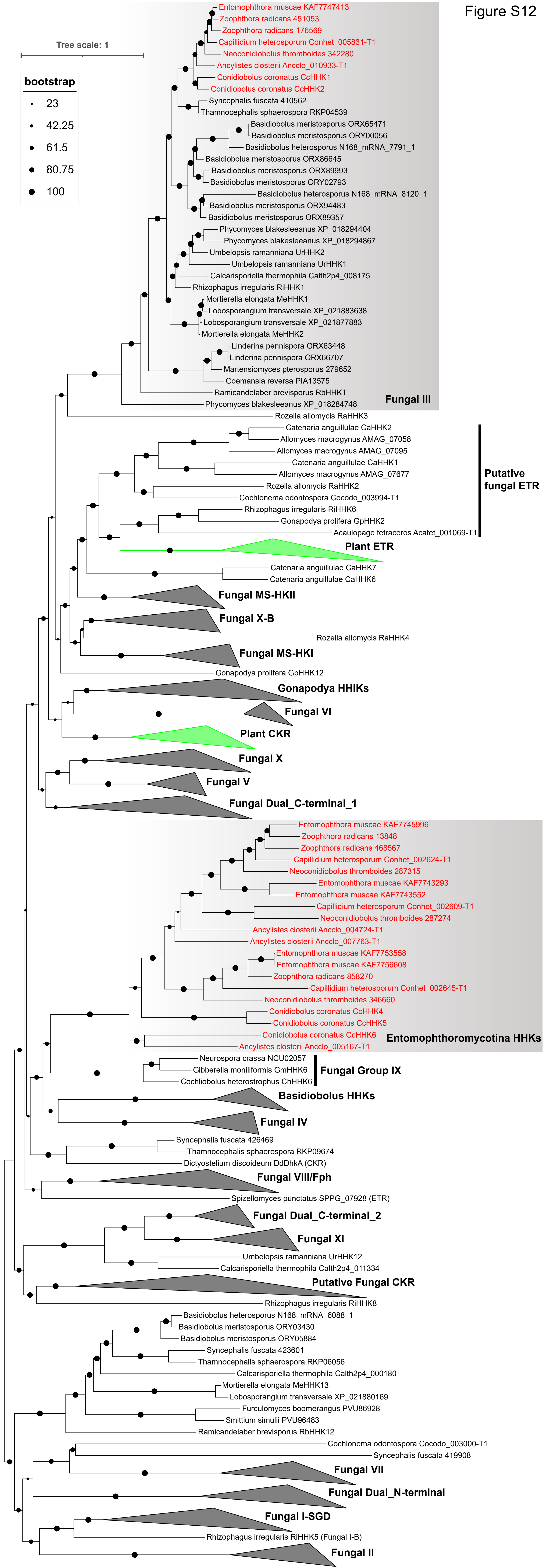

Tree scale: 1
bootstrap
23
42.25
61.5
80.75
100
Entomophthora muscae KAF7747413
Zoophthora radicans 451053
Zoophthora radicans 176569
Capillidium heterosporum Conhet_005831-T1
Neoconidiobolus thromboides 342280
Ancylistes closterii Ancclo_010933-T1
Conidiobolus coronatus CcHHK1
Conidiobolus coronatus CcHHK2
Syncephalis fuscata 410562
Thamnocephalis sphaerospora RKP04539
Basidiobolus meristosporus ORX65471
Basidiobolus meristosporus ORY00056
Basidiobolus heterosporus N168_mRNA_7791_1
Basidiobolus meristosporus ORX86645
Basidiobolus meristosporus ORX89993
Basidiobolus meristosporus ORY02793
Basidiobolus heterosporus N168_mRNA_8120_1
Basidiobolus meristosporus ORX94483
Basidiobolus meristosporus ORX89357
Phycomyces blakesleeanus XP_018294404
Phycomyces blakesleeanus XP_018294867
Umbelopsis ramanniana UrHHK2
Umbelopsis ramanniana UrHHK1
Calcarisporiella thermophila Calth2p4_008175
Rhizophagus irregularis RiHHK1
Mortierella elongata MeHHK1
Lobosporangium transversale XP_021883638
Lobosporangium transversale XP_021877883
Mortierella elongata MeHHK2
Linderina pennispora ORX63448
Linderina pennispora ORX66707
Martensiomyces pterosporus 279652
Coemansia reversa PIA13575
Ramicandelaber brevisporus RbHHK1
Phycomyces blakesleeanus XP_018284748
Fungal III
Rozella allomycis RaHHK3
Catenaria anguillulae CaHHK2
Allomyces macrogynus AMAG_07058
Allomyces macrogynus AMAG_07095
Catenaria anguillulae CaHHK1
Allomyces macrogynus AMAG_07677
Rozella allomycis RaHHK2
Cochlonema odontospora Cocodo_003994-T1
Rhizophagus irregularis RiHHK6
Gonapodya prolifera GpHHK2
Aculopage tetraceros Acatet_001069-T1
Putative fungal ETR
Plant ETR
Catenaria anguillulae CaHHK7
Catenaria anguillulae CaHHK6
Fungal MS-HKII
Fungal X-B
Rozella allomycis RaHHK4
Fungal MS-HKI
Gonapodya prolifera GpHHK12
Gonapodya HHIKs
Fungal VI
Plant CKR
Fungal X
Fungal V
Fungal Dual_C-terminal_1
Entomophthora muscae KAF7745996
Zoophthora radicans 13848
Zoophthora radicans 468567
Capillidium heterosporum Conhet_002624-T1
Neoconidiobolus thromboides 287315
Entomophthora muscae KAF7743293
Entomophthora muscae KAF7743552
Capillidium heterosporum Conhet_002609-T1
Neoconidiobolus thromboides 287274
Ancylistes closterii Ancclo_004724-T1
Ancylistes closterii Ancclo_007763-T1
Entomophthora muscae KAF7753558
Entomophthora muscae KAF7756608
Zoophthora radicans 858270
Capillidium heterosporum Conhet_002645-T1
Neoconidiobolus thromboides 346660
Conidiobolus coronatus CcHHK4
Conidiobolus coronatus CcHHK5
Conidiobolus coronatus CcHHK6
Ancylistes closterii Ancclo_005167-T1
Entomophthoromycotina HHKs
Neurospora crassa NCU02057
Gibberella moniliformis GmHHK6
Cochliobolus heterostrophus ChHHK6
Fungal Group IX
Basidiobolus HHKs
Fungal IV
Syncephalis fuscata 426469
Thamnocephalis sphaerospora RKP09674
Dictyostelium discoideum DdDhkA (CKR)
Fungal VIII/Fph
Spizellomyces punctatus SPPG_07928 (ETR)
Fungal Dual_C-terminal_2
Fungal XI
Umbelopsis ramanniana UrHHK12
Calcarisporiella thermophila Calth2p4_011334
Putative Fungal CKR
Rhizophagus irregularis RiHHK8
Basidiobolus heterosporus N168_mRNA_6088_1
Basidiobolus meristosporus ORY03430
Basidiobolus meristosporus ORY05884
Syncephalis fuscata 423601
Thamnocephalis sphaerospora RKP06056
Calcarisporiella thermophila Calth2p4_000180
Mortierella elongata MeHHK13
Lobosporangium transversale XP_021880169
Furculomyces boomerangus PVU86928
Smittium simulii PVU96483
Ramicandelaber brevisporus RbHHK12
Cochlonema odontospora Cocodo_003000-T1
Syncephalis fuscata 419908
Fungal VII
Fungal Dual_N-terminal
Fungal I-SGD
Rhizophagus irregularis RiHHK5 (Fungal I-B)
Fungal II
